# Trehalose metabolism and its impact on PrfA activity in *Listeria monocytogenes*

**DOI:** 10.64898/2026.05.05.722976

**Authors:** Jessica Schüler, Annette Walz, Niclas Wüstefeld, Maja Andiel, Wolfgang Eisenreich, Jeanine Rismondo

## Abstract

*Listeria monocytogenes* can grow as a saprophyte on decaying plant material, but can also switch to a pathogenic lifestyle. This switch is mediated by the virulence regulator PrfA, which activates the expression of most virulence genes. PrfA activity is tightly regulated by several mechanisms to ensure that virulence genes are only expressed within the host. One of these regulatory mechanisms is the sugar-dependent repression. In the presence of readily metabolizable sugars, which are imported via phosphotransferase systems (PTS) such as cellobiose, PrfA is repressed; however, the precise mechanism is still unknown. Using a sugar screen, trehalose was identified as the first PTS-dependent sugar that supports growth of *L. monocytogenes*, but does not seem to impact PrfA activity. We demonstrated that the PTS permease TreB is the sole trehalose importer. After import, trehalose-6-phosphate is cleaved by the phosphotrehalase TreA; however, loss of TreA does not fully abolish growth on trehalose suggesting that *L. monocytogenes* encodes an additional phosphotrehalase. ^13^C-Labeling experiments revealed that trehalose metabolism is repressed in the presence of glucose, while it can be metabolized in the presence of glycerol. Additionally, these experiments provided evidence that trehalose and cellobiose are metabolized via identical pathways, including glycolysis and the incomplete TCA cycle, although trehalose has a slower uptake and/or metabolization rate. We therefore hypothesize that sugar-dependent PrfA repression correlates with sugar transport and/or consumption rates, potentially due to varying availability of phosphoenolpyruvate (PEP), which serves as both a metabolic intermediate and phosphate donor for PTS-dependent transport.

## Importance

Virulence factors determine the pathogenic potential of bacteria; however, constant expression of these factors is an energetic burden. In *L. monocytogenes*, virulence gene expression is induced by the virulence regulator PrfA, whose activity is repressed in the presence of readily metabolizable sugars. This study reveals that trehalose, unlike other PTS-imported sugars, supports listerial growth while barely affecting PrfA activity, highlighting an exception to the established model of sugar-dependent PrfA repression. Based on ^13^C-Labeling experiments, we hypothesize that the metabolic flux rather than the sugar type alone influences PrfA activity. The proposed link between phosphoenolpyruvate availability and PrfA activity offers new insight into how nutrient sensing is linked to pathogenicity.

## Introduction

*Listeria monocytogenes* is a Gram-positive, rod-shaped bacterium belonging to the phylum *Bacillota.* It can alternate between a saprophytic and pathogenic lifestyle. In its saprophytic state, *L. monocytogenes* is ubiquitous in the environment, commonly found living on decaying plant material. Its resilience to various stressors such as elevated salt concentrations, acidic conditions, and temperature fluctuations, as well as nutrient limitation, allows *L. monocytogenes* to inhabit a variety of environments, including food industry, thereby increasing its pathogenic potential (1, 2). Through ingestion of contaminated food, namely dairy products, meat, seafood and vegetables, *L. monocytogenes* causes foodborne disease. Usually, infection with this bacterium is restricted to a self-limiting gastroenteritis in healthy individuals. However, infection in immunocompromised people causes severe invasive listeriosis or even neonatal infection in the case of pregnant women (3). Because of these factors, *L. monocytogenes* remains a significant concern for public health.

Due to its dual lifestyle, *L. monocytogenes* constantly needs to adapt its metabolism in response to the available nutrients and carbon sources in the environment and within the host (4). Hence, *L. monocytogenes* EGD-e possesses a range of ABC-transporters as well as the highest number of *pts* genes among bacterial species (5) encoding seven families of phosphoenolpyruvate (PEP)-dependent transport systems (PTS): PTS^Glc^, PTS^Man^, PTS^Lac^, PTS^Fru^, PTS^Gut^, PTS^Gat^ and PTS^Asc^. These permeases allow import of a variety of sugars with differing efficiency (6). Consequently, this enables *L. monocytogenes* to metabolize a wide variety of carbon sources including glucose, mannose, fructose and cellobiose (7, 8).

The switch from saprophytic growth in the environment to a pathogenic lifestyle is regulated by the master virulence regulator PrfA. PrfA is essential for survival within the host, as it activates and regulates the expression of key virulence genes in *L. monocytogenes* during infection. PrfA expression and activity itself underlie tight regulation including transcriptional, translational, and post-translational regulation mechanisms, to ensure that *L. monocytogenes* can respond to both environmental and host-derived signals (1, 4). Interestingly, PrfA activity is influenced by the type of available carbon source, known as sugar-dependent PrfA regulation. Environmental sugars such as cellobiose, glucose and fructose are imported via the PTS and subsequently enter glycolysis in the bacterial cell. These sugars are known to inhibit PrfA with cellobiose exerting the strongest inhibitory effect (4, 9–12), however, the precise mechanism of sugar-dependent PrfA repression is still not understood. It has been postulated that unphosphorylated EIIA components of PTS permeases, which accumulate in the presence of PTS-dependent sugars, interact and sequester PrfA thereby inhibiting its activity (1, 6). However, this protein-protein interaction has not been proven so far.

Using a sugar screen, we identified trehalose as the first PTS-dependent sugar which does not inhibit PrfA. Trehalose is a disaccharide composed of two α-D-glucose units linked by a α,α-1,1-glycosidic bond (13). TreB was previously identified as the trehalose-specific PTS permease in *L. monocytogenes* strain 1386 (14). We confirmed that TreB is also the sole trehalose transporter in the *L. monocytogenes* laboratory wildtype strain EGD-e. Interestingly, the phosphotrehalase TreA, which is required for the hydrolysis of trehalose-6-phosphate, is not essential for growth on trehalose as sole carbon source. In *Bacillus subtilis*, the expression of the *treBA* operon is controlled by the repressor TreR (15). We here confirm that TreR also acts as a repressor for the *treBA* operon in *L. monocytogenes*. Furthermore, we show that the *treBA* operon is highly expressed in complex media, while it is barely expressed in *Listeria* synthetic medium containing glucose as sole carbon source. Trehalose is, similar to cellobiose and glucose, metabolized via glycolysis; however, it is not consumed by *L. monocytogenes* in the presence of glucose, suggesting that trehalose metabolism is subject to carbon catabolite repression.

## Materials and methods

### Bacterial strains and growth conditions

All strains and plasmids used in this study are listed in Table S1. *Escherichia coli* strains were cultivated in lysogeny broth (LB) medium and *Listeria monocytogenes* strains in brain heart infusion (BHI) medium or *Listeria* synthetic medium (LSM) at 37°C unless otherwise stated. LSM was prepared as previously described using glucose, cellobiose or trehalose as sole carbon source (7). When required, media were supplemented with antibiotics at the following concentrations: for *E. coli* cultures: 30 µg ml^-1^ chloramphenicol (cat), 50 µg ml^-1^ kanamycin (kan), 100 µg ml^-1^ ampicillin (amp); for *L. monocytogenes* cultures: 7.5 µg ml^-1^ chloramphenicol, 50 µg ml^-1^ kanamycin, 5 µg ml^-1^ erythromycin (ery), 30 µg ml^-1^ nalidixic acid (nal).

### Strain and plasmid construction

All primers used in this study are listed in Table S2. For the markerless deletion of *manR* (*lmo0785*), *celR* (*lmo1721*) and *treA* (*lmo1254*), 1-kb DNA fragments up- and downstream of *manR, celR* and *treA* were amplified by PCR using primer pairs JR465/466 and JR467/468 (*manR*), JR471/472 and JR473/474 (*celR*) and JR494/495 and JR496/497 (*treA*), respectively. The resulting PCR products were fused in a second PCR using primer pair JR465/468 for *manR,* JR471/474 for *celR* and JR494/497 for *treA*. For the markerless inframe deletion of *treB* (*lmo1255*), *treR* (*lmo1253*) and *lmo1017*, 0.5-kb DNA fragments up- and downstream of *treB*, *treR* and *lmo1017* were amplified by PCR using primer pairs JR507/434 and JR435/508 (*treB*), JR511/501 and JR502/512 (*treR*) and NW11/12 and NW13/14 (*lmo1017*), respectively. The resulting PCR products were fused in a second PCR using primer pair JR507/508 for *treB*, JR511/512 for *treR* and NW11/NW14 for *lmo1017*. For the markerless inframe deletion of *lmo0184* and *lmo0862*, 0.7-kb DNA fragments up- and downstream of *lmo0184* and *lmo0862* were amplified by PCR using primer pairs JR563/564 and JR565/566 (*lmo0184*) and JS21/22 and JS23/24 (*lmo0862*), respectively. The resulting PCR products were fused in a second PCR using primers JR563/566 for *lmo0184* and JS21/24 for *lmo0862*. The *manR, celR, treA*, *treB*, *treR, lmo1017* and *lmo0184* deletion fragments were subsequently digested with *Kpn*I and *Sal*I and ligated into plasmid pKSV7 that had been cut with the same enzymes. The *lmo0862* deletion fragment was cut with *BamH*I and *Sal*I and ligated into *BamH*I/*Sal*I cut pKSV7. Plasmids pKSV7-Δ*manR*, pKSV7-Δ*celR*, pKSV7-Δ*treA*, pSKV7-Δ*treB*, pKSV7-Δ*treR,* pKSV7-Δ*lmo1017,* pKSV7-Δ*lmo0184* and pKSV7-Δ*lmo0862* were recovered in *E. coli* DH5α yielding strains EJR379, EJR378, EJR388, EJR389, EJR390, EJR404, EJR443 and EJR415, respectively. In addition, the plasmids pKSV7-Δ*manR*, pKSV7-Δ*celR,* pKSV7-Δ*treA*, pSKV7-Δ*treB*, pKSV7-Δ*treR,* pKSV7-Δ*lmo1017,* pKSV7-Δ*lmo0184* and pKSV7-Δ*lmo0862* were transformed into *L. monocytogenes* strain EGD-e and genes *manR*, *celR*, *treA*, *treB*, *treR, lmo1017, lmo0184* and *lmo0862* deleted by allelic exchange as previously described (16). The deletion of the respective genes was verified by PCR resulting in the construction of strains EGD-e Δ*manR* (LJR584), EGD-e Δ*celR* (LJR436), EGD-e Δ*treA* (LJR717), EGD-e Δ*treB* (LJR718), EGD-e Δ*treR* (LJR719), EGD-e Δ*lmo1017* (LJR746), EGD-e Δ*lmo0184* (LJR794) and EGD-e Δ*lmo0862* (LJR791). Plasmids pKSV7-Δ*lmo0184* and pKSV7-Δ*lmo0862* were also transformed into *L. monocytogenes* strain EGD-e Δ*treA* (LJR717) and genes *lmo0184* and *lmo0862* deleted by allelic exchange, resulting in the construction of strains EGD-e Δ*treA* Δ*lmo0184* (LJR793) and EGD-e Δ*treA* Δ*lmo0862* (LJR792).

For the constitutive overexpression of *treBA* and *treB*, *treBA* and *treB* were amplified by PCR using primer pairs JR491/492 and JR491/493, respectively. The resulting PCR products were cut with *Nco*I and *Sal*I and ligated into pIMK2 that had been cut with the same enzymes. Plasmids pIMK2-*treBA* and pIMK2-*treB* were recovered in *E. coli* DH5α yielding strains EJR383 and EJR385, respectively. The same plasmids were subsequently transformed into *L. monocytogenes* EGD-e yielding strains EGD-e pIMK2-*treBA* (LJR646) and EGD-e pIMK2-*treB* (LJR644). For the constitutive overexpression of *lmo0184* and *lmo0862*, both genes were amplified by PCR using primer pairs JR561/562 and JR551/552, respectively. The resulting PCR products were cut with *BamH*I and *Sal*I and ligated into pIMK2 that had been cut with the same enzymes. Plasmids pIMK2-*lmo0184* and pIMK2-*lmo0862* were recovered in *E. coli* DH5α yielding strains EJR444 and EJR417, respectively. Additionally, plasmids pIMK2-*lmo0184* and pIMK2-*lmo0862* were transformed into *L. monocytogenes* EGD-e and EGD-e Δ*treA* (LJR717) yielding strains EGD-e pIMK2-*lmo0184* (LJR776), EGD-e Δ*treA* pIMK2-*lmo0184* (LJR778), EGD-e pIMK2-*lmo0862* (LJR760) and EGD-e Δ*treA* pIMK2-*lmo0862* (LJR761).

For the construction of pPL3e-*P_actA_-lacZ* and pPL3e-*P_treBA_-lacZ*, 300 bp and 250 bp regions upstream of *actA* and *treB* were amplified using primer pairs JP3/JP4 and JR489/490, respectively. The resulting PCR products were digested with *BamH*I and *Sal*I and ligated into plasmid pPL3e-*lacZ* that had been cut with the same enzymes. Plasmids pPL3e-*P_actA_-lacZ* and pPL3e-*P_treBA_-lacZ* were recovered in *E. coli* DH5α yielding strains EJR286 and EJR382. In addition, plasmids pPL3e-*P_actA_-lacZ* and pPL3e-*P_treBA_-lacZ* were transformed into *L. monocytogenes* EGD-e yielding strains EGD-e pPL3e-*P_actA_-lacZ* (LJR352) and EGD-e pPL3e-*P_treBA_-lacZ* (LJR622). For the construction of EGD-e Δ*treR* pPL3e-*P_treBA_-lacZ* (LJR730), plasmid pPL3e-*P_treBA_-lacZ* was first transformed into *E. coli* strain S17-1 yielding strain EJR403 and subsequently transferred into EGD-e Δ*treR* via conjugation.

For overproduction of His-TreR in *E. coli*, plasmid pWH844-*treR* was constructed. For this purpose, the *treR* gene was amplified using primers NW21 and NW22. The resulting PCR product was digested with *BamH*I and *Sal*I and ligated into plasmid pWH844 that had been cut with the same enzymes. pWH844-*treR* was subsequently recovered in *E. coli* DH5α yielding strain EJR410.

### Lecithinase (PlcB) activity assay

For a qualitative PlcB activity assay, single colonies of the indicated *L. monocytogenes* strains were streaked on LB agar supplemented with 0.2% activated charcoal and 2% of an egg yolk suspension. The egg yolk suspension was prepared by mixing one egg yolk with an equal volume of 1x phosphate-buffered saline (PBS, pH 7.4). To assess the sugar-dependent PrfA repression, 25 mM cellobiose or 25 mM trehalose were additionally added to the plates. Inoculated plates were incubated for 24 hours at 37°C and pictures were taken.

### Growth curves

For growth assays, the indicated *L. monocytogenes* strains were grown overnight in BHI broth or LSM glucose at 37°C with shaking. The next morning, overnight cultures were diluted in fresh BHI broth or LSM glucose to an OD_600_ of 0.1 and grown at 37°C and 200 rpm until an OD_600_ of 0.3 was reached. Cells of 2 ml culture were collected per condition, washed twice with 1 ml of the corresponding LSM and re-suspended in 1 ml LSM. The cultures were adjusted to an OD_600_ of 0.1 and 200 µl of the cell suspension transferred into wells of a 96-well plate. To compare the growth of the *L. monocytogenes* wildtype strain EGD-e in LB and LB containing 25 mM trehalose, LB overnight cultures of EGD-e were diluted to an OD_600_ of 0.1 in fresh LB medium and grown until an OD_600_ of 0.3 was reached. Cells of 2 ml culture were collected, washed twice in LB broth or LB broth containing 25 mM trehalose. The pellets were subsequently re-suspended in 1 ml of the respective medium and cultures adjusted to an OD_600_ of 0.1. 200 µl of the cell suspension were transferred into wells of a 96-well plate. For all growth assays, plates were incubated at 37°C with orbital shaking in the BioTek Epoch 2 microplate reader and the OD_600_ was measured every 15 min for at least 25 hours. Averages and standard deviations of at least three independent growth assays were plotted.

### Expression and purification of His-TreR

For the overproduction of His-TreR, plasmid pWH844-*treR* was transformed into *E. coli* strain DH5α. The resulting strain was grown in LB supplemented with Amp at 37°C until an OD_600_ of 0.8 – 1.0 was reached. The expression of *his*-*treR* was induced by the addition of 1 mM isopropyl β-D-1-thiogalactopyranoside (IPTG). The culture was incubated for another 1.5 h at 37°C. Cells were subsequently harvested by centrifugation at 4,330 x g for 15 min, washed once with 1x cell disruption buffer (ZAP; 50 mM Tris-HCl, pH 7.5, 200 mM NaCl) and the cell pellet stored at -20°C until further use. The cell pellet was resuspended in 15 ml 1x ZAP buffer and the cells were passaged three times through an HTU DIGI-F press (18,000 lbf/in^2^; G. Heinemannn, Schwäbisch Gmünd, Germany). Afterwards, the cell debris was collected by centrifugation at 13,600 x g and 4°C for 15 min, followed by a second centrifugation of the supernatant at 126,000 x g and 4°C for 45 min. The supernatant was subjected to a Ni^2+^ nitrilotriacetic acid column (IBA, Göttingen, Germany) and His-TreR eluted using an imidazole gradient. Elution fractions were subsequently analyzed by SDS-PAGE and selected fractions dialyzed against 1x ZAP buffer at 4°C overnight. Protein concentrations were determined by Bradford protein assay (17) using the Bio-Rad protein assay dye reagent concentrate. For the standard curve, bovine serum albumin was used. The protein samples were either directly used or snap frozen and stored at -80°C until further use.

### Electrophoretic mobility shift assay

236- and 200-bp DNA fragments containing the *treBA* and *cadA* promoter were amplified using primers NW23/24 and NW25/26, respectively. To determine the binding ability of His-TreR to the *treBA* promoter, 20 µl EMSA reactions were prepared. These contained 0.5, 1, 1.5 and 2 µmol of His-TreR, 100 ng *treBA* promoter DNA and 2 µl of 10x EMSA buffer (100 mM Tris base, 200 mM NaCl, 2 mM Na_2_EDTA, pH 8.0). A reaction of 2 µmol of His-TreR with 100 ng *cadA* promoter DNA served as negative control. EMSA reactions were incubated for 20 min at room temperature. After addition of 4 µl of 6x EMSA loading dye (0.2% bromophenol blue, 0.2% xylene cyanole in 50% glycerol), the samples were separated on 8% native acrylamide gels (8% acrylamide, 1x EDTA-free Tris-borate-EDTA (TBE) buffer, pH 8.0 [100 mM Tris base, 50 mM boric acid, 12.5 mM NaCl], 0.08% ammonium persulfate, 0.08% tetramethylethylenediamine) in 0.5x EMSA buffer. A pre-run of the gels was performed for 30 min at 90 V before the samples were loaded. The run was performed at 4°C for 30 min at 80 V followed by 1.5 h at 100 V. The gels were stained in 50 ml 0.5x EMSA buffer containing 5 µl HDGreen Plus DNA dye (INTAS, Göttingen, Germany) for 30 sec. The DNA bands were visualized using a Gel Doc XR+ (Bio Rad, Munich, Germany).

### β-Galactosidase assay

For the comparison of the *P_treBA_* promoter activity in different media, strain EGD-e pPL3e-*P_treBA_-lacZ* (LJR622) was grown as follows: 1) overnight cultures were grown in 10 ml LSM glucose containing 5 µg ml^-1^ erythromycin, diluted to an OD_600_ of 0.1 in 15 ml LSM glucose; 2) overnight cultures were grown in 10 ml BHI broth containing 5 µg ml^-1^ erythromycin, diluted to an OD_600_ of 0.1 in 15 ml BHI broth; 3) overnight cultures were grown in 10 ml BHI broth containing 5 µg ml^-1^ erythromycin, washed twice in (a) LSM glucose (b) LSM trehalose and cultures adjusted to an OD_600_ of 0.1 in the respective LSM; 4) overnight cultures were grown in 10 ml LB broth containing 5 µg ml^-1^ erythromycin, diluted to an OD_600_ of 0.1 in 15 ml LB broth; 5) overnight cultures were grown in 10 ml LB broth containing 5 µg ml^-1^ erythromycin, washed twice in (a) LSM glucose (b) LSM trehalose and cultures adjusted to an OD_600_ of 0.1 in the respective LSM. For the comparison of the *P_actA_* promoter activity in different media, strain EGD-e pPL3e-*P_actA_-lacZ* (LJR352) was grown as follows: 1) overnight cultures were grown in 10 ml BHI broth containing 5 µg ml^-1^ erythromycin, diluted to an OD_600_ of 0.1 in 15 ml BHI broth containing 1% amberlite; 2) overnight cultures were grown in 10 ml LB broth containing 5 µg ml^-1^ erythromycin, diluted to an OD_600_ of 0.1 in 15 ml LB broth containing 1% amberlite. Where indicated, BHI and LB amberlite cultures contained additionally 25 mM glucose, cellobiose or trehalose. 3) overnight cultures were grown in 10 ml LSM glucose containing 5 µg ml^-1^ erythromycin, washed twice in (a) LSM glucose (b) LSM cellobiose (c) LSM trehalose and cultures adjusted to an OD_600_ of 0.1 in the respective LSM; 4) overnight cultures were grown in 10 ml LSM glucose with low BCAA concentrations containing 5 µg ml^-1^ erythromycin, washed twice in (a) LSM glucose (b) LSM cellobiose (c) LSM trehalose with low BCAA concentrations and cultures adjusted to an OD_600_ of 0.1 in the respective LSM. Cultures were grown at 37°C and 200 rpm until they reached an OD_600_ of 0.6-0.8. The final OD_600_ of each culture was measured prior sample collection. 1 ml of each culture was collected, re-suspended in 100 µl assay buffer with Triton X-100 (ABT buffer) (60 mM K_2_HPO_4_, 40 mM KH_2_PO_4_, 100 mM NaCl, 0.1% Triton X-100; pH 7), snap-frozen in liquid nitrogen, and stored at -80°C until further use. The β-galactosidase assay was performed as described previously (18). Briefly, samples were thawed, and 10-fold dilutions were prepared in ABT buffer. 50 µl of each dilution were mixed with 10 µl of 0.4 mg ml^-1^ 4-methyl-umbelliferyl-β-D-galactopyranoside (MUG) substrate (Merck, Darmstadt, Germany) that was dissolved in dimethyl sulfoxide and incubated for 60 min in the dark. Reactions containing ABT buffer and the substrate were used as blank. Next, 20 µl of each reaction were transferred into the wells of a black 96-well plate containing 180 µl ABT buffer. Fluorescence values were determined using a Synergy Mx microplate reader (BioTek) at 366 nm excitation and 445 nm emission wavelengths. Concentrations from 0.015625 to 4 µM of the fluorescent standard 4-methylumbelliferone (MU) were used to obtain a standard curve. β-galactosidase units, or MUG units, were calculated as (pmol of substrate hydrolyzed x dilution factor)/(culture volume in ml x OD_600_ x reaction time in min). The amount of hydrolyzed substrate was determined from the standard curve as (emission reading – *y* intercept)/slope).

### ^13^C-Labeling experiment with [U-^13^C_6_]glucose in the presence of cellobiose and trehalose

One single colony of *L. monocytogenes* EGD-e was picked from agar plates and transferred into 50 ml LB medium. Three of these 50 ml EGD-e cultures were incubated overnight at 37°C and 200 rpm and united after growing to one culture. Three 200 ml main cultures were adjusted to an OD_600_ of 0.1 by adding 29 ml of EGD-e overnight culture into LB medium, which were treated with different carbon sources. To each EGD-e main culture, [U-^13^C_6_]glucose (99% ^13^C-abundance, Sigma-Aldrich) was added to a final concentration of 2 mM. Additionally, a final concentration of 25 mM trehalose or 25 mM cellobiose (natural ^13^C-abundance) was added, while no additional carbon source was added to the third culture. The EGD-e main cultures were grown to an OD_600_ of 0.5-0.6. A volume of 2 ml of 2 M sodium azide was added to kill the *L. monocytogenes* EGD-e cells. The suspension was centrifuged at 4,000 x g (Megafuge 2.0R, Heraeus instruments) for 15 min at 22°C. The pellets were washed three times with 1 ml phosphate-buffered saline (PBS, pH 7.4) by centrifugation at 4,000 x g (Megafuge 2.0R, Heraeus instruments) for 5 min at 22°C. Afterwards, the pellets were stored in the fridge. The experiment was performed in triplicate.

### ^13^C-Labeling experiment with [U-^13^C_3_]glycerol in the presence of cellobiose and trehalose

To eliminate glucose from the medium of the main cultures, the following experiment was performed by using LSM containing 1% glycerol. For this purpose, 50 ml LSM containing 1% glucose were inoculated with a single colony of *L. monocytogenes* EGD-e. Four of these 50 ml EGD-e cultures were incubated overnight at 37°C and 200 rpm. The cultures were washed two times by centrifugation [22°C, 10 min, 4,000 x g (Megafuge 2.0R, Heraeus instruments)] with 7.5 - 10 ml LSM containing 1% glycerol to remove the glucose-containing medium. Four 200 ml main cultures were adjusted to an OD_600_ of 0.1 by adding 7.5 - 10 ml of EGD-e cells into LSM containing 1% unlabelled glycerol, which were subsequently treated with different carbon sources. To three EGD-e main cultures, 30 µl of [U-^13^C_3_]glycerol (99% ^13^C abundance, Sigma-Aldrich) was added to a final concentration of 2 mM. Two cultures additionally contained either a final concentration of 25 mM trehalose or 25 mM cellobiose (natural ^13^C abundance). As a control, one 200 ml main culture was only treated with 2.5% unlabelled glycerol. All EGD-e main cultures were grown to an OD_600_ of 0.4-0.5. 1 ml sodium azide was added to a final concentration of 10 mM to kill the *L. monocytogenes* EGD-e cells. The suspension was centrifuged at 4,000 x g (Megafuge 2.0R, Heraeus instruments) for 15 min at 22°C. The pellets were washed three times with 2 ml PBS (pH 7.4) by centrifugation at 4,000 x g (Megafuge 2.0R, Heraeus instruments) for 15 min at 22°C. Afterwards, the pellets were stored in the fridge. The experiment was performed in triplicate.

### Acidic hydrolysis of cell mass and derivatisation of resulted amino acids from *L. monocytogenes* strain EGD-e

*L. monocytogenes* EGD-e cell pellets were freeze-dried for 24 h. 2 mg of pellet (dry weight) were hydrolyzed with 6 M HCl for 24 h at 105°C. After cooling down, the solvent was removed under a stream of nitrogen gas at 70°C. The residues were dissolved in 200 µl 70% acetic acid. Amino acids were purified by column chromatography (Dowex 50WX8, H^+^-form, 37 - 74 µm, about 0.3 ml). The ion exchange column was washed with 70% methanol and water. After the sample had been transferred to the column, the column was washed with water. The elution was performed with aqueous 4 M NH_3_. The eluate was collected, and the solvent was removed under a stream of nitrogen gas at 70°C. For derivatization, 50 µl acetonitrile and 50 µl *N*-(*tert*-butyldimethylsilyl)-*N*-methyl-trifluoroacetamide containing 1% *tert*-butyldimethylsilylchloride were added to the residue and incubated for 30 min at 70°C.

### Isolation of cell wall monosaccharides and lipids from *L. monocytogenes* strain EGD-e labelled with [U-^13^C_6_]glucose

10 g of freeze-dried pellet was incubated in 0.5 ml 3 M MeOH/HCl at 80°C overnight. The supernatant was collected, and the solvent was removed under a stream of nitrogen gas at room temperature. For derivatization, the residue was incubated in 1 ml acetone containing 20 µl conc. H_2_SO_4_ at room temperature for 1 h. 2 ml saturated NaCl and 2 ml saturated Na_2_CO_3_ solutions were added to the sample. The mixture was extracted twice with 3 ml ethyl acetate. The organic phases were collected, and the volume was reduced under reduced pressure at 40°C. The solvent was removed under a stream of nitrogen gas at room temperature. After dissolving the residue in 200 µl anhydrous ethyl acetate: acetic anhydride solution (1:1), the mixture was heated up to 60°C overnight. Afterwards, the solvent was dried under a stream of nitrogen gas at room temperature. The residue, containing diisopropylidene/acetate derivates, was dissolved in 100 µl anhydrous ethyl acetate and studied by gas chromatography/mass spectrometry (GC/MS) analytics.

### Isolation of cell wall monosaccharides and lipids from *L. monocytogenes* strain EGD-e labelled with [U-^13^C_3_]glycerol

10 g of the freeze-dried pellet was hydrolyzed and methylated in 0.5 ml 3 M MeOH/HCl for 6 h at 80°C. The solvent was removed under a stream of nitrogen gas at room temperature. For derivatization, the residue was incubated in 1 ml acetone containing 20 µl conc. H_2_SO_4_, at room temperature for 1 h. 2 ml saturated NaCl and 2 ml saturated Na_2_CO_3_ solutions were added to the sample. The mixture was extracted two times with 3 ml ethyl acetate. The organic phases were collected, and the solvent was removed under a stream of nitrogen gas at room temperature. After dissolving the residue in 200 µl anhydrous ethyl acetate: acetic anhydride solution (1:1), the mixture was heated up at 60 °C overnight. Afterwards, the solvent was dried under a stream of nitrogen gas at room temperature. The residue, containing diisopropylidene/acetate derivates, was dissolved in 100 µl anhydrous ethyl acetate and studied by GC/MS analytics.

### Gas chromatography/mass spectrometry analytics (GC/MS)

GC/MS analytic was performed with a QP2010 Plus gas chromatography/mass spectrometry (Shimadzu), linked with a Silica capillary column (Equity TM-5; 30 x 0.25 mm, 0.25 µm film thickness, SUPELCO). An electron impact ionization at 70 eV and a quadrupole detector were used. Each sample was measured three times to obtain three technical replicates. Additionally, three biological replicates of each sample were analysed. The software GC/MS solution software (Shimadzu) was used for data acquisition.

### Calculation of the ^13^C-excess and isotopologue profiles

The calculation of ^13^C-excess and isotopologue abundances was carried out by an in-house Excel-based macro employing the solver function of Excel. For this method, the natural occurrence of the ^13^C-isotope was determined by measuring standards (compounds without additional ^13^C-enrichment). In addition, *de novo* synthesized compounds in the presence of ^13^C-labeled carbon sources were measured. After collecting all necessary data, the linear regression analysis, which can be divided into 3 basic steps, was performed: (i) Determination of the impact of derivatization reagent was determined by calculated theoretical abundance (in a matrix) of the measured molecule, which was compared to natural abundances of standard. The calculated multiplication vector determined the input of the derivatization reagent. (ii) In a second matrix, values of the ^13^C-enriched sample were represented in a vector. The multiplication vector was normalized and corresponded to absolute enrichment (natural + non-natural ^13^C-abundance). (iii) The matrix is constructed like in (i). The resulting multiplication vector was normalized to represent ^13^C-excess values (19–21). Data of ^13^C-labelled compounds with an total excess of at least 2% after growth with [U-^13^C_6_]glucose are shown. For consistency, the same compounds were chosen for the [U-^13^C_3_]glycerol experiments. The total excess of the control condition, namely growth in LB with [U-^13^C_6_]glucose and LSM with [U-^13^C_3_]glycerol, were set to 1. The mean and standard deviations of normalized total excess and isotopologue profiles were plotted for selected amino acids, cell wall sugars and fatty acids.

## Results

### PlcB activity is not affected by trehalose

The activity of the main virulence regulator PrfA of *L. monocytogenes* is strongly influenced by the available carbon sources (4); however, the precise mechanism of sugar-dependent PrfA repression is still not fully understood. So far, sugar-dependent PrfA repression was only assessed for a subset of carbon sources that can be metabolized by *L. monocytogenes*, such as glucose or cellobiose (6, 9–11, 22, 23). Thus, a screen of a variety of sugars was performed to determine their impact on PrfA activity (data not shown). As a read-out of PrfA activity, PlcB activity assays were used, which rely on the ability of PlcB to cleave lecithin in egg yolk, resulting in a characteristic white halo around the bacterial colonies. Indeed, white halos can be observed for the *L. monocytogenes* wildtype and the *prfA* complementation strain, when they were grown on LB plates containing activated charcoal and egg yolk (Fig. 1). In accordance with previous studies, halos were absent when *prfA* is deleted or in the presence of cellobiose (Fig. 1)(10, 24). Interestingly, PlcB activity can still be observed for the *L. monocytogenes* wildtype and the *prfA* complementation strain in the presence of trehalose (Fig. 1). To ensure that *L. monocytogenes* metabolizes trehalose when grown in LB medium, growth experiments were performed in the absence and presence of trehalose. The *L. monocytogenes* wildtype strain EGD-e showed enhanced growth in the presence of the disaccharide, confirming its utilization under these conditions (Fig. S1).

**Fig. 1:**
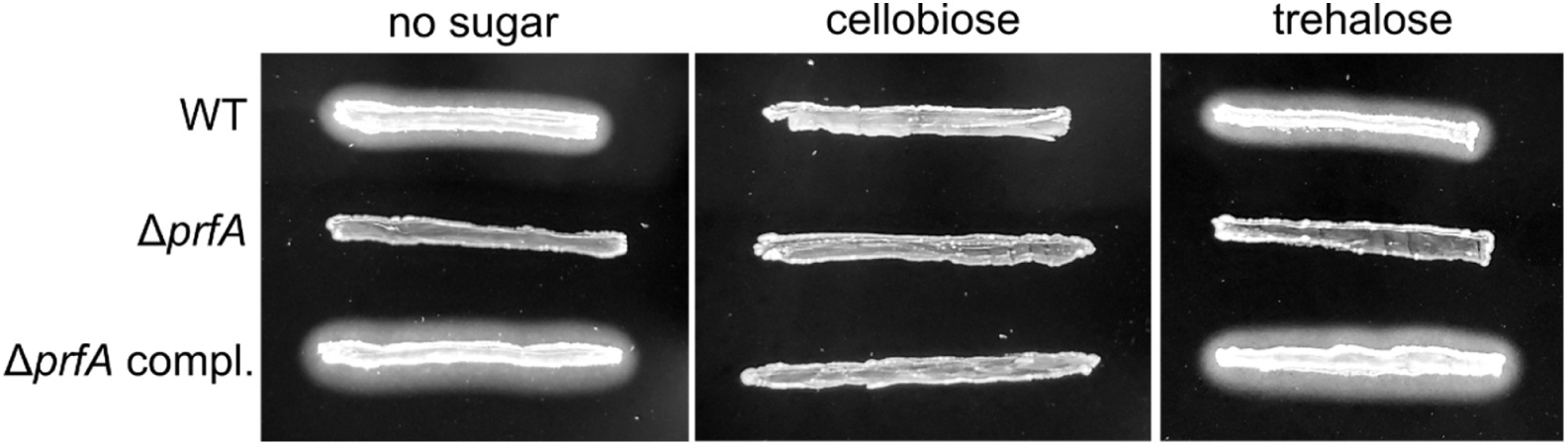
PlcB activity in the presence of different carbon sources. PlcB activity of the *L. monocytogenes* wildtype strain EGD-e (WT), the *prfA* mutant and the *prfA* complementation strain was tested on LB agar plates containing activated charcoal and egg yolk or on plates that additionally contained 25 mM cellobiose or trehalose. A representative image of three independent experiments is shown.

### Lmo1071, TreA and TreB are required for growth on trehalose

Trehalose is thought to be solely imported by the PTS-permease TreB, consisting of an EIIB and EIIC domain (14). After import, trehalose is cleaved into glucose and glucose-6-phosphate by the phosphotrehalase TreA (25). Based on homology, it was proposed that TreR acts as a repressor for the *treBA* operon (15). To verify the involvement of TreA, TreB and TreR in trehalose metabolism in the *L. monocytogenes* wildtype strain EGD-e, deletion mutants of the respective genes were constructed. No growth differences in LSM with glucose or cellobiose as sole carbon source were observed between the wildtype and the *treA*, *treB* and *treR* deletion mutants (Fig. S2A-B). Interestingly, growth of the wildtype strain was slightly reduced when it was shifted from LSM glucose to LSM trehalose (Fig. 2B) in comparison to the shift from BHI to LSM trehalose (Fig. 2A). In contrast, growth of the *treR* deletion was comparable between both pre-growth conditions (Fig. 2A-B). In the absence of *treB*, *L. monocytogenes* is unable to grow in LSM with trehalose as sole carbon source (Fig. 2A-B), which is in accordance with previous studies (14). TreB is the PTS permease consisting of EIIB and EIIC domains; however, the corresponding EIIA protein involved in trehalose metabolism is currently unknown. It was previously hypothesized that TreB is phosphorylated by the glucose/glucosidase class EIIA encoded by *lmo1017* (12). We thus tested the impact of Lmo1017 on the growth of *L. monocytogenes* in LSM trehalose. The deletion of *lmo1017* led to a minor growth defect, when the strain was shifted from BHI broth to LSM trehalose (Fig. 2C). This growth defect was more pronounced when the strain was shifted from LSM glucose to LSM trehalose (Fig. 2D), suggesting that Lmo1017 is indeed involved in trehalose metabolism. Interestingly, the *treA* deletion strain had only a minor growth deficit in LSM trehalose, when the strain was pre-grown in BHI (Fig. 2A). In contrast, when the *treA* mutant was pre-grown in LSM glucose, the strain only started to grow in LSM trehalose after approximately 15 hours (Fig. 2B). These results suggest that *L. monocytogenes* encodes another enzyme that can cleave trehalose-6-phosphate. Two potential candidates were identified by a blast search with the TreA protein of *L. monocytogenes* as query: Lmo0184 and Lmo0862, which share 43% identity and 61% similarity and 36% identity and 55% similarity with TreA, respectively. If one or both of these proteins are able to cleave trehalose-6-phosphate, their overexpression should improve the growth of a *treA* deletion strain in LSM trehalose. In contrast, deletion of *lmo0184* or *lmo0862* should reduce the ability of the *treA* mutant to grow in LSM trehalose after it was pre-grown in BHI broth. However, neither overexpression nor deletion of *lmo0184* and *lmo0862* affected the growth of the *treA* mutant in LSM trehalose (Fig. S3). Interestingly, overproduction of *lmo0184* led to a slight growth defect of both, the wildtype and the *treA* mutant, in LSM glucose (Fig. S3A). These results suggest that neither Lmo0184 nor Lmo0862 are involved in trehalose metabolism and that *L. monocytogenes* encodes another, so far unknown enzyme that is able to cleave trehalose-6-phosphate.

**Fig. 2:**
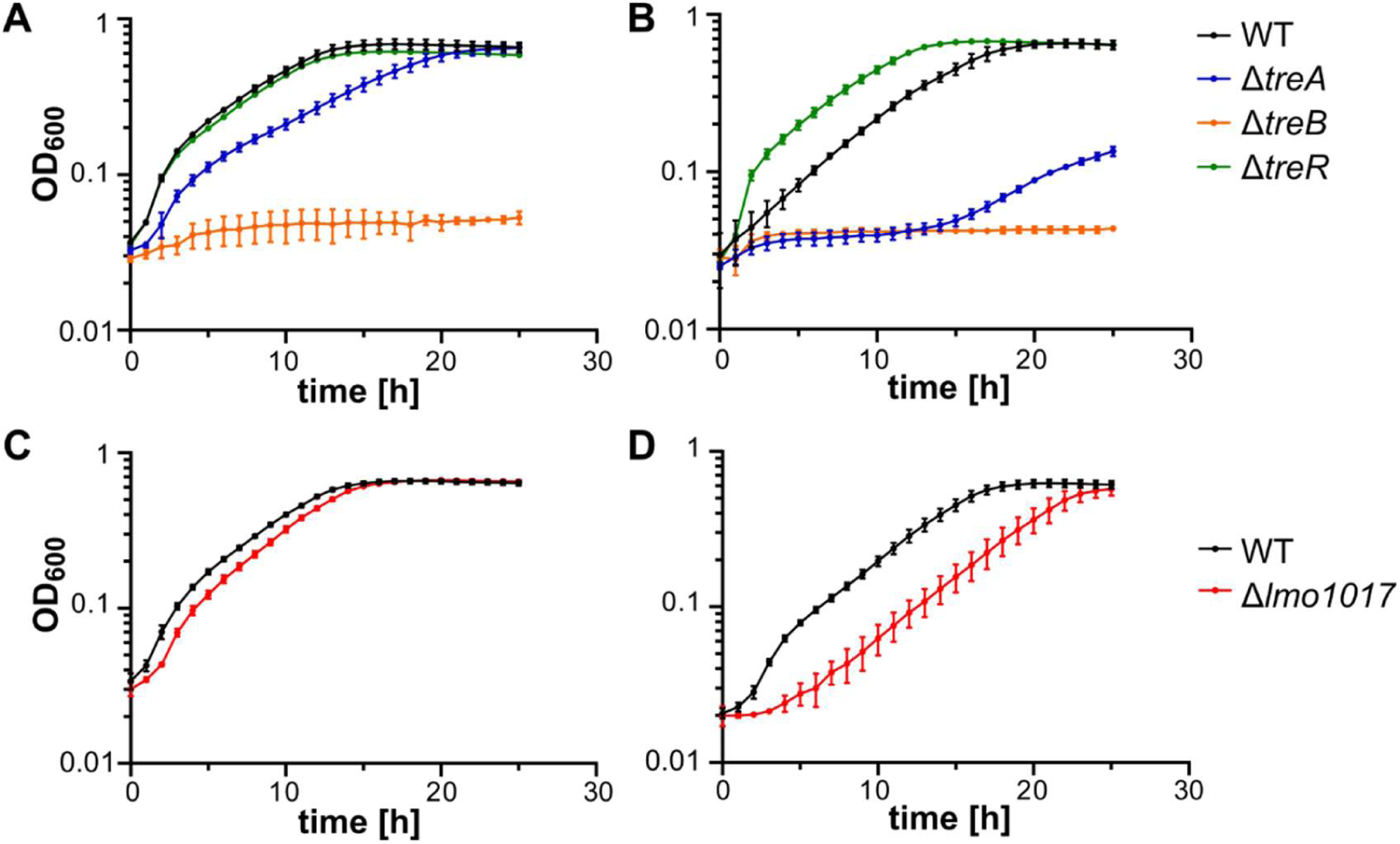
Growth of *treA*, *treB*, *treR* and *lmo1017* deletion strains on LSM trehalose. (A, C) The indicated *L. monocytogenes* strains were grown overnight in BHI broth, diluted to an OD_600_ of 0.1 in fresh BHI broth and grown until an OD_600_ of 0.3 was reached. (B, D) *L. monocytogenes* strains were grown overnight in LSM glucose, diluted to an OD_600_ of 0.1 in fresh LSM glucose and grown until an OD_600_ of 0.3 was reached. (A-D) Cells were collected, washed with LSM trehalose and OD_600_ adjusted as described in the methods section. The growth was monitored for 25 hours using a plate reader. The average values and standard deviations of the OD_600_ readings of three independent experiments were plotted.

### Overproduction of TreB leads to a growth defect

Next, we wanted to determine whether the overexpression of the *treBA* operon, which might lead to an enhanced trehalose metabolism, has an impact on PrfA activity. However, no differences in PlcB activity were observed between the wildtype strain, a *treB* and a *treBA* overexpression strain (Fig. S4A). Furthermore, no growth difference was observed between the wildtype strain and the *treBA* overexpression strain in LSM glucose or LSM trehalose. Interestingly, overproduction of the PTS permease TreB alone led to a growth defect in LSM trehalose, while growth was similar to the wildtype strain in LSM glucose (Fig. S4B), likely due to the accumulation of trehalose-6-phosphate.

### The *treBA* operon is highly expressed in LB medium and LSM trehalose

Growth of the *L. monocytogenes* wildtype strain EGD-e in LSM trehalose seems to be influenced by the pre-growth condition (Fig. 2). Indeed, promoter activity assays revealed that the activity of the *P_treBA_* promoter is highly dependent on the growth condition. Medium activity levels were observed when *L. monocytogenes* is grown in BHI medium, while high promoter activity was measured for bacteria that were grown in LSM trehalose or LB medium (Fig. 3A, C). In contrast, the *treBA* operon was barely expressed when *L. monocytogenes* was grown in LSM glucose (Fig. 3B). Interestingly, the *P_treBA_* promoter activity was lowered when the bacteria were shifted from BHI or LB medium to LSM glucose (Fig. 3A, C). As indicated above, TreR is proposed to regulate the expression of the *treBA* operon (15). To test whether TreR is able to bind to the promoter region of the *treBA* operon, an EMSA assay was performed. Indeed, *P_treBA_* and TreR form a complex resulting in a shift of the DNA band in the native gel. In contrast, no complex formation was observed for EMSA reactions containing TreR and the promoter of *cadA*, a reaction that served as negative control (Fig. S5). To determine whether TreR indeed acts as a repressor, *P_treBA_* activity was compared between the wildtype strain and the *treR* deletion strain. In the absence of *treR*, no changes between the *P_treBA_* activity can be observed between *L. monocytogenes* that had been grown in BHI medium or that had been shifted from BHI medium to LSM glucose (Fig. 3A). In addition, the *P_treBA_* activity was higher for the *treR* mutant that had been grown in LSM glucose as compared to the wildtype strain, confirming that TreR acts as a repressor of the *treBA* operon.

**Fig. 3:**
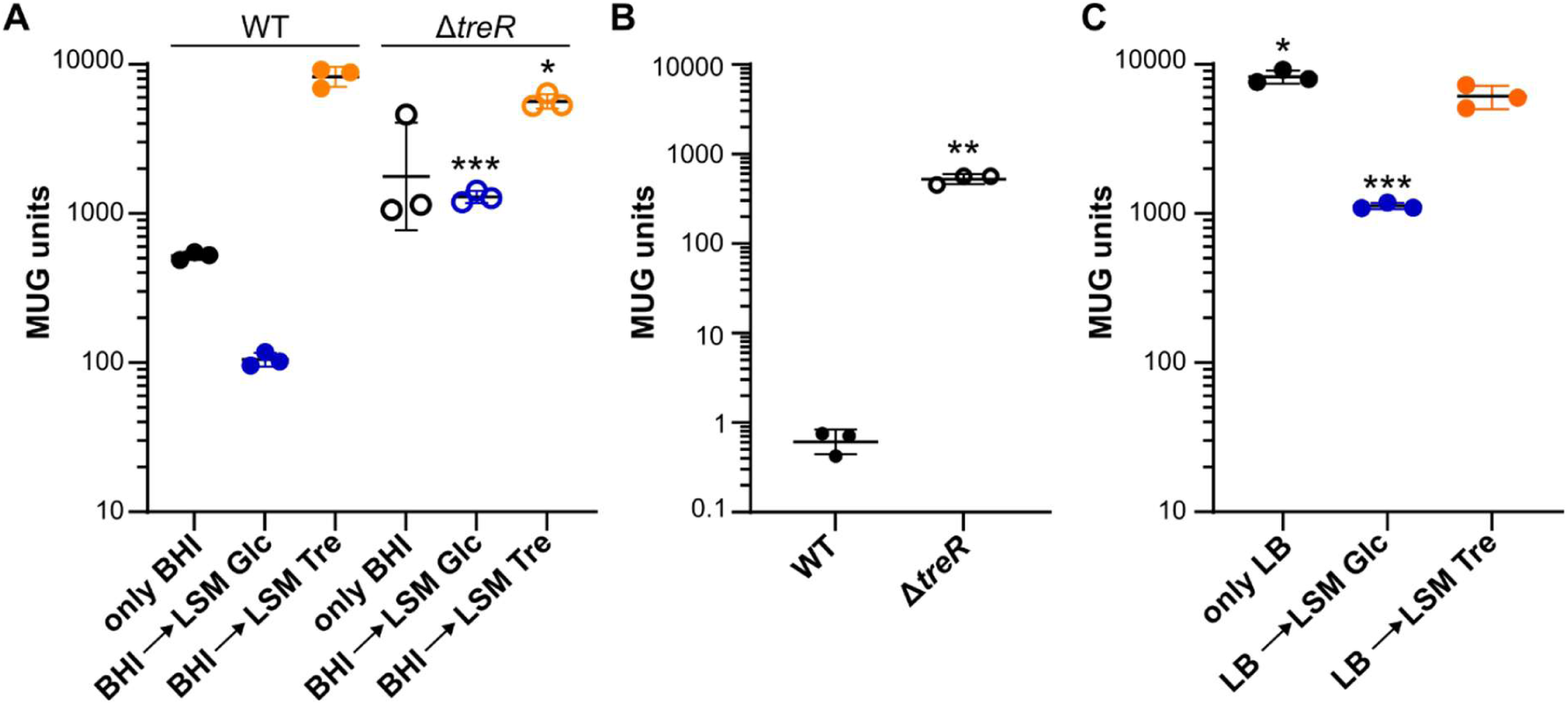
The *treBA* operon is differentially expressed in diverse media. *L. monocytogenes* strains EGD-e pPL3e-*P_treBA_-lacZ* and EGD-e Δ*treR* pPL3e-*P_treBA_-lacZ* were grown in diverse media as described in the methods section. For panel C, *P_treBA_* activity was only tested for EGD-e pPL3e-*P_treBA_-lacZ*. The activity of the *treBA* promoter activity was subsequently determined by performing β-galactosidase activity assays. The averages of the β-galactosidase activity and standard deviations of three independent experiments were plotted. For data presented in panel A and B, statistical significance was determined using an unpaired t-test with Welch’s correction comparing wildtype and the *treR* mutant. For statistical analysis of data presented in panel C, a one-way ANOVA coupled with a Dunnett’s multiple comparison test using the growth condition LB ◊ LSM trehalose as reference was used (*, *p* ≤ 0.5; **, *p* ≤ 0.01, ***, *p* ≤ 0.001).

### Catabolite repression of trehalose uptake by [U-^13^C_6_]glucose

Among the PTS carbohydrates identified to date, cellobiose exerts the strongest repression of PrfA activity in *L. monocytogenes* (12, 24). It is therefore even more surprising that the PTS transported sugar trehalose, which differs from cellobiose only in its glycosidic bond, did not inhibit PrfA in our assays. This observation led to the hypothesis that the use of different metabolic pathways could be responsible for the distinct regulatory effects of these carbohydrates. To validate this, a retrosynthetic analysis using [U-^13^C_6_]glucose and [U-^13^C_3_]glycerol in the presence of either unlabeled cellobiose or unlabeled trehalose was performed. In line with previous labeling experiments with *L. monocytogenes*, an efficient uptake of [U-^13^C_6_]glucose from the medium was observed, resulting in a high total excess in several metabolites (26, 27). In particular, significant labelling was found in the amino acids aspartate (Asp; 20.1% ± 0.5%), alanine (Ala; 12.5% ± 0.3%), cell wall-bound sugars such as ribose (26.6% ± 1.2%) and galactose (53.2%± 3.5%), and the fatty acids C15:0 (23.3% ± 2.5%), C17:0 (24.1% ± 2.3%)(Fig. 4A, C, E). Supplementation of unlabeled cellobiose in the medium resulted in a significant reduction in total excess in Asp (11.13% ± 0.19%), Ala (6.76% ± 0.12%), C15:0 (16.6% ± 6.1%), C17:0 (15.4% ± 3.2%), ribose (18.7% ± 0.7%) and galactose (37.6% ± 0.5%). The total excess values of amino acids glutamate (Glu; 4.7% ± 0.4%), lysine (Lys; 2.1% ± 0.13%) and serine (Ser; 2.1% ± 0.3%) were also reduced in the presence of cellobiose to 2.6% ± 0.17%, 1.1% ± 0.2% and 1.5% ± 0.1%, respectively. In addition to the proteinogenic amino acids, high total excess values were obtained for diaminopimelic acid (DAP; 45.3% ± 1.3%). The presence of cellobiose reduced the total excess of DAP to 25.4% ± 0.3% (Fig. 4A). These data confirm an active metabolism and cellular uptake of cellobiose in EGD-e in the presence of [U-^13^C_6_]glucose. Interestingly, no dilution effects were observed when unlabeled trehalose was added to the medium as no reduction of the total excess was detected in *de novo* synthesized amino acids, fatty acids or cell wall compounds (Fig. 4A, C, E). This provided clear evidence for catabolite repression of trehalose uptake and metabolism in the presence of [U-^13^C_6_]glucose. In contrast, trehalose utilization was observed when *L. monocytogenes* was cultivated with [U-^13^C_3_]glycerol, which was measurable by a reduced total ^13^C excess in all analyzed metabolites in the presence of trehalose (Fig. 5A, C, E). Together, these results indicated that [U-^13^C_6_]glucose mediated catabolite repression of trehalose uptake, while under the same conditions, cellobiose utilization was unaffected.

**Fig. 4:**
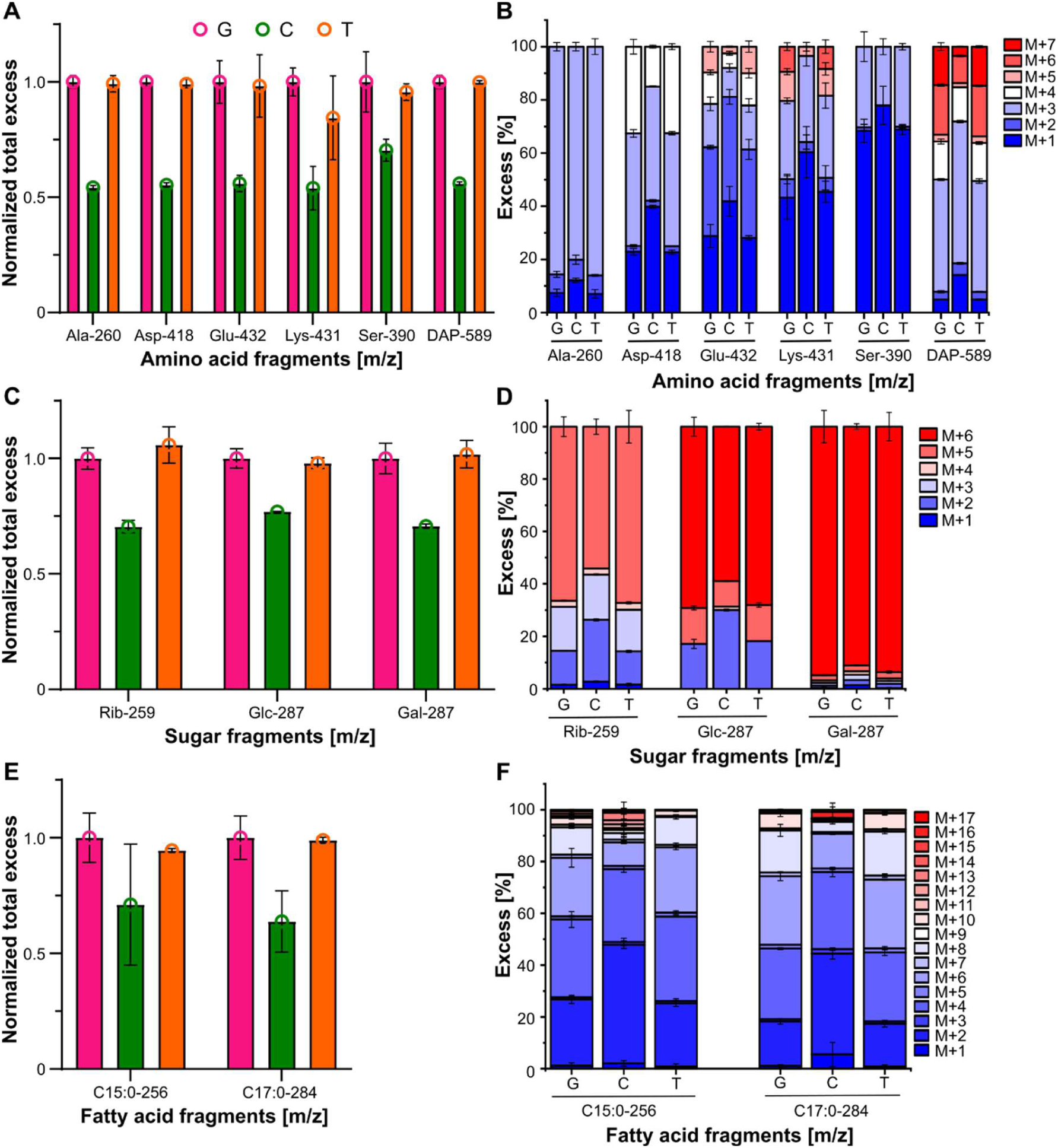
Isotopologue profiling of amino acids, cell wall sugars and fatty acids of *L. monocytogenes* cultivated in the presence of [U-^13^C_6_]glucose. (A, C, E) Normalized ^13^C-excess (total excess) and (B, D, F) ^13^C-isotopologue fractions (normalized to 100%) of (A-B) amino acids, (C-D) cell wall sugars and (E-F) fatty acids of *L. monocytogenes* grown in LB broth in the presence of [U-^13^C_6_]glucose. LB broth was additionally supplemented with 25 mM cellobiose (label C) or 25 mM trehalose (label T). LB broth without additional carbon sources was used as reference (label G). Mean and standard deviations of three biological replicates are plotted.

**Fig. 5:**
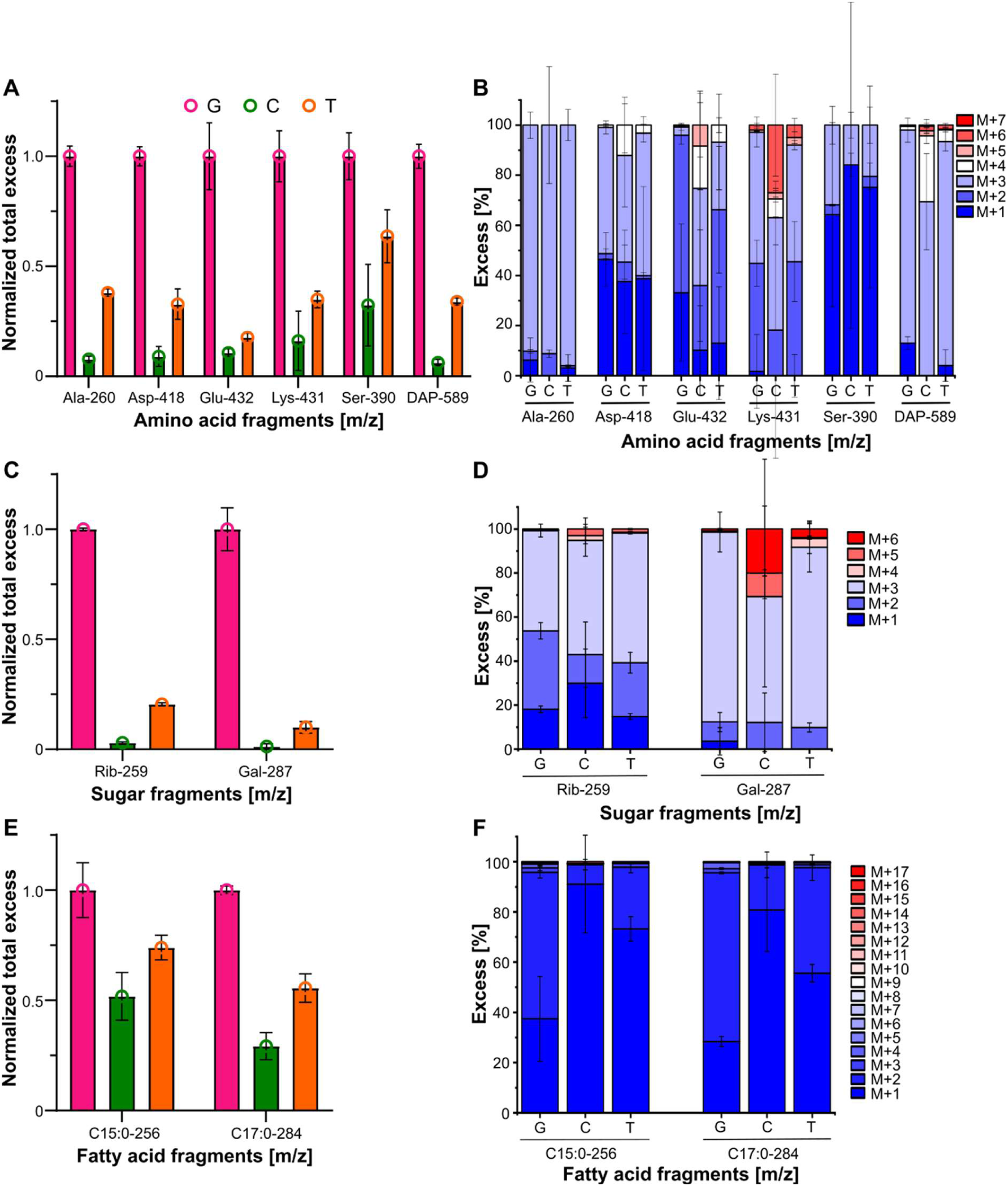
Isotopologue profiling of amino acids, cell wall sugars and fatty acids of *L. monocytogenes* cultivated in the presence of [U-^13^C_3_]glycerol. (A, C, E) Normalized ^13^C-excess (total excess) and (B, D, F) ^13^C-isotopologue fractions (normalized to 100%) of (A-B) amino acids, (C-D) cell wall sugars and (E-F) fatty acids of *L. monocytogenes* grown in LB broth in the presence of [U-^13^C_3_]glycerol. LSM was additionally supplemented with 25 mM cellobiose (label C) or 25 mM trehalose (label T). LSM without additional carbon sources was used as reference (label G). Mean and standard deviations of three biological replicates are plotted.

To obtain detailed information about the carbon flux distribution of cellobiose, isotopologue profiles of *L. monocytogenes* cultivated on labeled [U-^13^C_6_]glucose were evaluated. In particular, the labeling patterns of newly synthesized compounds, such as amino acids, fatty acids, and cell wall bound sugars allowed to reconstruct the pathways from the ^13^C_6_-precursor into the products. The amino acid fragment Ala-260 (m/z) predominantly showed M+3 (i.e. an isotopologue carrying three ^13^C-atoms), with minor contributions from M+1 and M+2, in the presence of [U-^13^C_6_]glucose (Fig. 4B). This indicated biosynthesis mainly from [U-^13^C_3_]pyruvate generated from [U-^13^C_6_]glucose via [U-^13^C_6_]glucose-6-phosphate through glycolysis, with a small amount from ^13^CO_2_ via carboxylation reactions and from unlabeled carbon pools. Cellobiose metabolism significantly reduced the percentage of the M+3 species by half (absolute mean value 5.45%) without altering the overall isotopologue pattern. The isotopologues M+1, M+3, and M+4 in Asp-418, in the presence of [U-^13^C_6_]glucose (Fig. 4B), confirmed its biosynthesis via the transamination of oxaloacetate, which was formed from glycolytic intermediates and the incomplete tricarboxylic acid (TCA) cycle. The uptake and metabolism of cellobiose led to a marked decrease in the M+3 and M+4 isotopologues, revealing metabolic fluxes of cellobiose into glycolysis and the incomplete TCA cycle. The reduction in the higher isotopologues in Glu-432 (m/z) further confirmed that cellobiose was degraded via the incomplete TCA cycle (Fig. 4B). The values for the isotopologue M+1 of Ser-390 (m/z) were similar between the different growth conditions, while the presence of cellobiose led to a reduced excess of M+2 compared to the culture cultivated only in the presence of [U-^13^C_6_]glucose and the culture cultivated in the presence of [U-^13^C_6_]glucose and trehalose. The uptake of cellobiose resulted in a strong reduction of the M+6 isotopologue of Lys-431 (m/z) in comparison to [U-^13^C_6_]glucose alone. Furthermore, the presence of cellobiose led to a reduced excess of M+4, M+5, M+6 and M+7 of DAP compared to cultivation of EGD-e with only [U-^13^C_6_]glucose. DAP is an intermediate in lysine biosynthesis (28). It is formed out of [U-^13^C_3_] or [U-^13^C_4_]aspartate and [U-^13^C_3_]pyruvate. The isotopologue profile showed a dominant M+3 isotopologue when only [U-^13^C_6_]glucose was present, which originates from [U-^13^C_3_]pyruvate or [U-^13^C_3_]aspartate. The combination of both fully labelled precursors led to the M+7 isotopomer (Fig. 4B).

Isotopic analysis of cell wall bound sugars and fatty acids provided insights into the gluconeogenesis, the pentose phosphate pathway, and fatty acid biosynthesis. In particular, glucose (Glc-287 (m/z)) and galactose (Gal-287 (m/z)) showed a predominant M+6 isotopologue caused by the fully labelled [U-^13^C_6_]glucose. For Gal-287 (m/z), only a small amount of the lower isotopologues was detected (Fig. 4D). This indicated an efficient conversion from [U-^13^C_6_]glucose into [U-^13^C_6_]galactose. The presence of cellobiose reduced the amount of the M+6 isotopologue in Gal-287 (m/z) and Glc-287 (m/z). The M+3 and M+5 isotopologues in ribose confirmed the biosynthesis via the pentose phosphate pathway in *L. monocytogenes* grown in the presence of [U-^13^C_6_]glucose. The metabolization of cellobiose diluted the M+5 isotopologue with a simultaneous increase of the M+2 isotopologue in the profile of ribose (Rib-259)(Fig. 4D). *L. monocytogenes* characteristic cell membrane fatty acids C15:0 and C17:0 predominantly exhibited M+2 or multiples of M+2, consistent with biosynthesis via the condensation of [U-^13^C_2_]acetyl-CoA with malonyl-CoA. Only a small fraction of fully ^13^C labeled fatty acid was detected. The dilution effect, caused by the metabolization of cellobiose, was detected in both fatty acids, measurable in the form of a reduction of higher isotopologues by a simultaneous increase of the M+2 isotopologue (Fig. 4F).

To summarize, across all analyzed compounds, cellobiose metabolism reduced the ^13^C enrichments and decreased corresponding isotopologues, but did not significantly alter the overall isotopologue pattern compared to cultures grown exclusively in [U-^13^C_6_]glucose. These data indicated that unlabelled cellobiose followed the same metabolic routes as the ^13^C-labelled glucose does, i.e. through glycolysis, an incomplete TCA cycle, the pentose phosphate pathway, fatty acid biosynthesis, and gluconeogenesis. Consequently, the ^13^C-excess values due to the incorporation of [U-^13^C_6_]glucose were decreased due to the dilution of ^13^C-labelled metabolic intermediates with unlabelled specimens derived from the unlabelled cellobiose supplement, but the isotopologue patterns (i.e. the relative fractions of ^13^C-isotopologues) remained unaffected. In contrast, when unlabelled trehalose was supplied together with [U-^13^C_6_]glucose, no changes or reductions in the total excess or in the isotopologue profiles were observed in EGD-e cultures across all analyzed metabolites (Fig. 4), demonstrating persistent catabolite repression.

### Trehalose and cellobiose were metabolized through identical metabolic pathways

Unlike glucose, glycerol is taken up independently of the PTS system through the GlpF aquaporin transporter (29). Furthermore, trehalose metabolism is not subject to catabolite repression in the presence of glycerol. The use of glycerol as a ^13^C-labelled carbon source thus enabled the characterization of trehalose flux contributions in *L. monocytogenes*; however, it is noteworthy that low isotopic enrichment increased variability in isotopologue profiles across biological replicates. This is especially pronounced for samples of *L. monocytogenes* grown in presence of [U-^13^C_3_]glycerol and cellobiose, which led to a marked reduction of ^13^C-labelled compounds.

As described earlier (27), the imported [U-^13^C_3_]glycerol is phosphorylated to glycerol-3-phosphate and dihydroxyacetone phosphate, allowing it to directly enter glycolysis and providing a well-defined starting point for carbon flux from the labelled source into central metabolism. In the presence of [U-^13^C_3_]glycerol, Ala-260 (m/z) exhibited a high fraction of the M+3 isotopologue in *L. monocytogenes*, indicating the inclusion of the [U-^13^C_3_]pyruvate label into Ala by transamination(Fig. 5B). Trehalose and cellobiose metabolism decreased the absolute amount of the M+3 fragment but did not influence the relative isotopologue patterns. A small M+4 fragment was found for Asp-418 (m/z) in the presence of all carbon sources. The M+3 fragment of Asp-418 showed the highest relative excess, followed by the M+1 fragment in the presence of [U-^13^C_3_]glycerol and trehalose. The isotopologue composition of Glu-432 (m/z) displayed characteristic M+1 and M+2 fractions (Fig. 5B), reflecting synthesis from [U-^13^C_2_]acetyl-CoA and the incomplete TCA cycle. However, the small fraction of ^13^C-glutamate (overall ^13^C-excess, 0.4% ± 0.06%) suggested an additional uptake of unlabeled glutamate or a related compound from the medium besides *de novo* biosynthesis of glutamate. The presence of cellobiose and trehalose led to a strong reduction of M+2 of DAP-589.

Under all conditions, the cell wall bound and galactose (Gal-287 (m/z)) showed dominant M+3 isotopologues (Fig. 5D) indicating synthesis from [U-¹³C₃]pyruvate via gluconeogenesis. A minor M+6 fraction in galactose determined a limited formation using two fully labeled pyruvate units. Ribose (Rib-259 (m/z)) also exhibited a high M+3 species (Fig. 5D), consistent with its biosynthesis via the pentose phosphate pathway from [^13^C_6_]glucose, generated through gluconeogenesis. The metabolism of cellobiose and trehalose diluted the M+3 pools in Gal-287 (m/z), and Rib-259 (m/z) (Fig. 5D).

The fatty acid C17:0 had a pronounced M+1 isotopologue (absolute mean value: 2.3% ± 0.16%), which increased to 2.75% ± 0.5% (absolute mean value) and 3.05% ± 0.2% (absolute mean value) when cellobiose and trehalose were taken up into the cell (Fig. 5F). The M+1 isotopologue was most likely formed by the incorporation of individual ^13^C-atoms from carbohydrate metabolism into acetyl-CoA or malonyl-CoA, which occurs via partially labeled intermediates or through CO_2_ fixation. The M+2 and M+4 isotopologues, although at low levels, reflected the incorporation of fully labeled acetyl-CoA and malonyl-CoA units during fatty acid biosynthesis. Both PTS sugars, cellobiose and trehalose, significantly reduced the M+2 and M+2 multimers, indicating metabolic fluxes through fatty acid biosynthesis under these conditions (Fig. 5F).

In sum, the simultaneous availability of cellobiose or trehalose in the presence of [U-^13^C_3_]glycerol did not result in any qualitative changes in the observed isotopologue patterns across all analysed metabolites in *L. monocytogenes* (Fig. 5). These data proved that both disaccharides, cellobiose and trehalose, entered central metabolism through the same routes and were metabolized via identical pathways, including glycolysis, the incomplete TCA, the pentose phosphate pathway, fatty acid biosynthesis, and gluconeogenesis.

### Trehalose was metabolized or imported more slowly than cellobiose

In cultures grown with [U-^13^C_3_]glycerol as the sole carbon source, small total excess values were detected for Ala-260 (m/z) (1.14% ± 0.05), Asp-418 (m/z) (1.11% ± 0.04%), Lys-431 (m/z) (0.29% ± 0.03%), Ser-390 (m/z) (0.27% ± 0.03%) and DAP-589 (m/z) (1.03% ± 0.06%). The uptake and metabolism of cellobiose reduced the total excess of Ala-260 (m/z), Asp-428 (m/z), Glu-432 (m/z), Lys-431 (m/z), Ser-390 (m/z) and DAP-589 (m/z) to 0.09% ± 0.02%, 0.10% ± 0.05%, 0.04% ± 0.01%, 0.05% ± 0.04%, 0.09% ± 0.05%, and 0.06% ± 0.01%, respectively. In contrast, trehalose only caused a moderate reduction of the total excess compared to the presence of only [U-^13^C_3_]glycerol, with remaining total excess values of 0.43% ± 0.02% for Ala-260 (m/z), 0.36% ± 0.08% for Asp-418 (m/z), 0.07% ± 0.01% for Glu-432 (m/z), 0.1% ± 0.01% for Lys-431 (m/z), 0.17% ± 0.03% for Ser-390 (m/z) and 0.35% ± 0.02% for DAP-589 (Fig. 5A). Consistent with the amino acid labelling profiles, the weaker reduction of total excess through the metabolization of trehalose in comparison to cellobiose was also evident in fatty acids C15:0 and C17:0 as well as the cell wall bound sugars ribose, and galactose (Fig. 5C, E). Together, these data suggest that trehalose was metabolized more slowly and/or imported through the PTS system at a lower rate than cellobiose. These results raised the question whether the slowed trehalose metabolism is responsible for the absence of trehalose-dependent PrfA repression.

### Reduction in sugar metabolism reduces repressive effects on PrfA

If the assumption is correct that PrfA is not repressed by trehalose due to a slow transport and/or metabolization of this sugar, then a reduction in the transport or metabolism of other sugars should likewise attenuate or abolish their repressive effects on PrfA. ManR and CelR act as transcriptional activators and control the expression of mannose- and glucose-specific PTS and cellobiose-specific PTS, respectively (11, 24, 30, 31). Their deletion thus likely leads to a reduced import and consumption of these sugars. Indeed, the growth of the *celR* deletion strain is slightly reduced in LSM containing cellobiose as sole carbon source. However, no growth differences were observed for the *manR* deletion strain, when grown in LSM glucose. Interestingly, growth of the *manR* and *treR* deletion strains was enhanced in LSM trehalose compared to the wildtype strain (Fig. S6). PlcB assays revealed that reduced cellobiose uptake in fact reduces its repressing effect on PrfA activity, as the *celR* deletion strain exhibits halo formation in the presence of cellobiose (Fig. 6). Strikingly, also the deletion of *manR* leads to PlcB activity in the presence of glucose (Fig. 6), although no growth difference was measured in LSM glucose (Fig. S6).

**Fig. 6:**
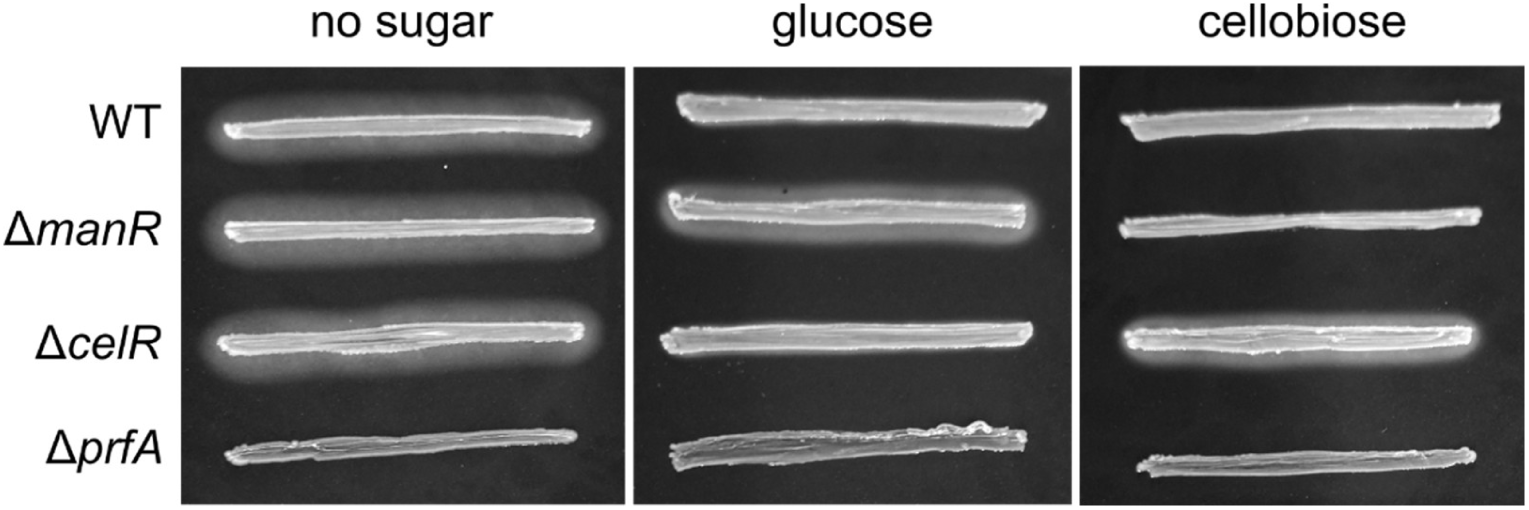
PlcB activity of *manR* and *celR* deletion strains. PlcB activity of the *L. monocytogenes* wildtype strain EGD-e (WT) and the *manR*, *celR* and *treR* mutants was tested on LB agar plates containing activated charcoal and egg yolk or on plates that additionally contained 25 mM glucose or cellobiose. A representative image of three independent experiments is shown.

### Trehalose-dependent PrfA repression varies across growth conditions

It has been shown frequently that PrfA activity varies depending on the culture medium and the available carbon source. We thus wanted to determine the PrfA activity in a variety of growth media, which are supplemented with either glucose, cellobiose and trehalose. For this purpose, the promoter of *actA*, which is a PrfA regulated gene, was fused to a promoter-less *lacZ* gene and integrated into the chromosome of *L. monocytogenes*. This strain was subsequently grown under diverse conditions and the promoter activity determined by β-galactosidase assays, which can be used as a read-out for PrfA activity. Indeed, these assays revealed strong differences in PrfA activity depending on the type of growth medium and the available carbon source (Fig. 7). Growth in BHI medium supplemented with amberlite, a PrfA activating agent, resulted in weak PrfA levels. The addition of 25 mM glucose or trehalose led to a minor decrease of PrfA activity, while PrfA activity was significantly attenuated in the presence of cellobiose (Fig. 7A). Surprisingly, all three carbon sources, glucose, cellobiose and trehalose, have a repressive effect on PrfA when *L. monocytogenes* is grown in LB amberlite (Fig. 7B). PrfA activity is also regulated by the global metabolic regulator CodY. High levels of branched-chain amino acids lead to inactivation of PrfA, while its activity is increased in low BCAA environments (32). We thus wanted to confirm that this regulation also occurs when *L. monocytogenes* is grown in LSM. Additionally, we wanted to determine the impact of glucose, cellobiose and trehalose on PrfA activity, when they are present as sole carbon source. In accordance with previous observations, PrfA activity is 10-fold higher when *L. monocytogenes* is grown in LSM with low BCAA levels compared to LSM with normal BCAA levels (Fig. 7C-D). Cellobiose exerted the strongest repressive effect on PrfA, regardless of the BCAA concentration (Fig. 7C-D). Surprisingly, PrfA inhibition was only observed in the presence of trehalose, when LSM was supplemented with low levels of BCAA (Fig. 7C-D). These results suggest that trehalose-dependent repression of PrfA strongly depends on the growth medium.

**Fig. 7:**
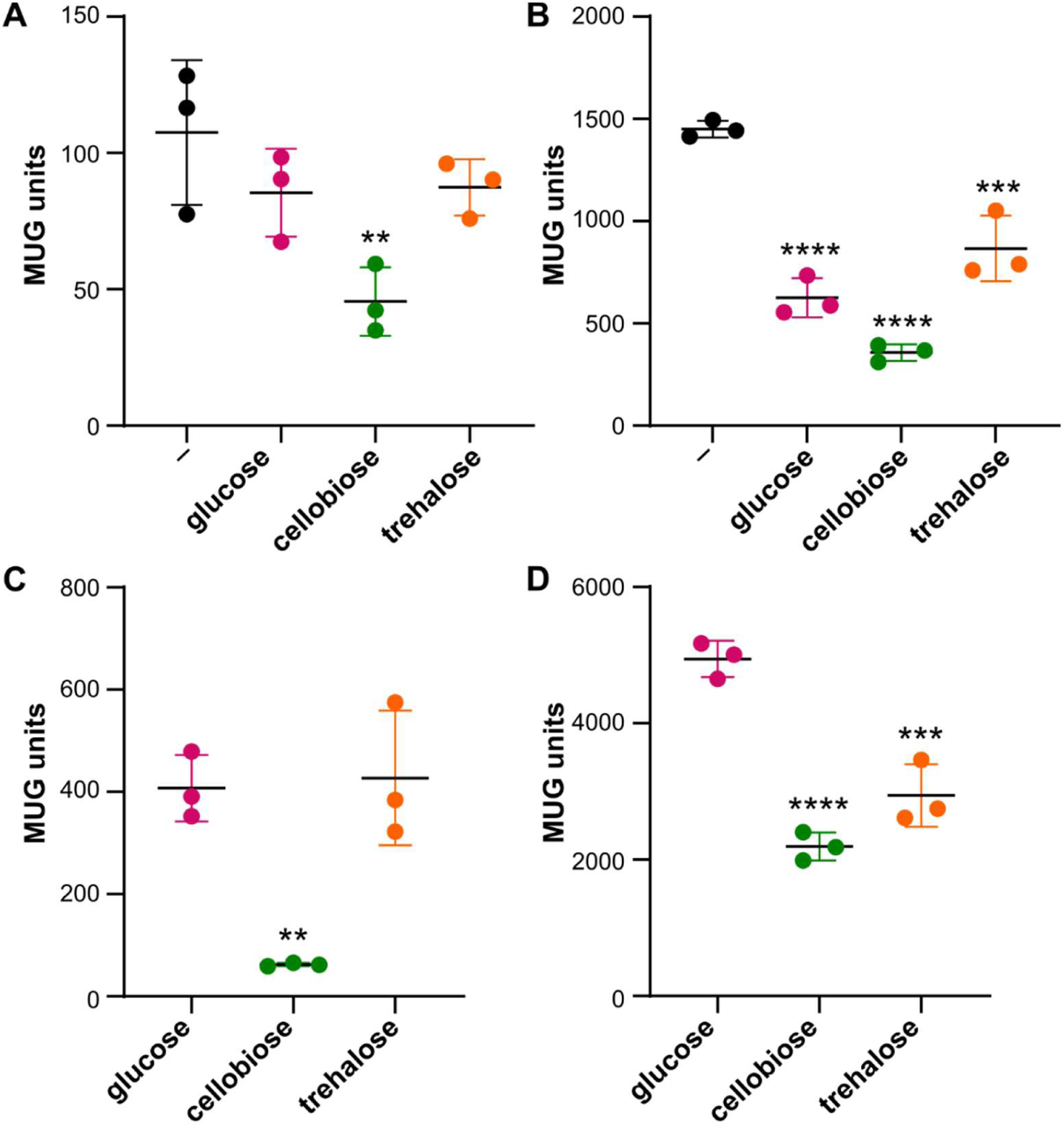
Trehalose-dependent PrfA repression depends on the medium. *L. monocytogenes* strain EGD-e pPL3e-*P_actA_-lacZ* was grown in (A) BHI containing 1% amberlite and additionally 25 mM of the indicated sugars, (B) LB containing 1% amberlite and additionally 25 mM of the indicated sugars, (C) LSM with glucose, cellobiose or trehalose as sole carbon source and (D) LSM with low BCAA levels and glucose, cellobiose or trehalose as sole carbon source as described in the methods section. The activity of the *treBA* promoter activity was subsequently determined by performing β-galactosidase activity assays. The averages of the β-galactosidase activity and standard deviations of three independent experiments were plotted. For statistical analysis, a one-way ANOVA coupled with a Dunnett’s multiple comparison test using (A-B) the growth condition without additional sugars (-) and (C-D) the LSM glucose growth condition as reference was used (**, *p* ≤ 0.01, ***, *p* ≤ 0.001, ****, *p* ≤ 0.0001).

## Discussion

*L. monocytogenes* can easily switch from a saprophytic to a pathogenic lifestyle. This switch is facilitated by the activity of the transcriptional regulator PrfA, which controls the expression of most virulence genes. PrfA itself is regulated on a transcriptional, translational, and post-translational level to ensure that virulence genes are only expressed when the bacterium enters its host (4). One of these regulation mechanisms is the sugar-dependent PrfA repression, which was identified more than three decades ago (10); however, it is still not fully understood. According to the current model, PrfA activity is repressed by readily metabolizable PTS-dependent sugars. It was further suggested that the glucose-dependent repression of PrfA is mediated by a phosphorylation of PrfA via ManL, an EIIA-EIIB component of PTS^Man^, although this phosphorylation was not experimentally proven (33). Previous studies mainly focused on the impact of glucose or cellobiose on PrfA; however, additional carbon sources that are present in the environment can support growth of *L. monocytogenes* (7). Thus, a sugar screen was used to determine whether other carbon sources also affect the activity of PrfA. Surprisingly, *L. monocytogenes* still showed PlcB activity in the presence of trehalose, suggesting that PrfA activity is not repressed under this condition. A previous study suggested that TreB is the sole PTS transporter in *L. monocytogenes* strain 1386 (14). Lack of TreB also led to an inability of the *L. monocytogenes* strain EGD-e to grow, when trehalose was used as a sole carbon source, confirming that trehalose import solely depends on this PTS permease. TreB consists of an EIIB and EIIC domain; however, no EIIA component for trehalose metabolism has been identified in *L. monocytogenes* so far. Deutscher and colleagues have previously suggested that *lmo1017* might encode an EIIA component that phosphorylates TreB (12). Here we show that deletion of *lmo1017* indeed resulted in a growth defect in LSM trehalose. However, growth was not completely abolished, suggesting that Lmo1017 as well as other EIIA proteins are involved in trehalose uptake. Similar observations were also made for *B. subtilis*, where three EIIA proteins, PtsA, PtsG and GamP, are able to support growth on trehalose (34).

TreB is encoded in an operon with the phosphotrehalase TreA, which hydrolyses trehalose-6-phosphate into glucose-6-phosphate and glucose (Fig. 8) (25). Interestingly, deletion of *treA* did not completely abolish the ability of *L. monocytogenes* to grow on trehalose as sole carbon source, suggesting that this organism encodes at least one additional enzyme with phosphotrehalase activity. The two closest homologs of TreA, Lmo0184 and Lmo0862, do not seem to be involved in trehalose metabolism. Thus, future research is required to identify an enzyme that is able to cleave trehalose-6-phosphate in the absence of TreA in *L. monocytogenes*.

**Fig. 8:**
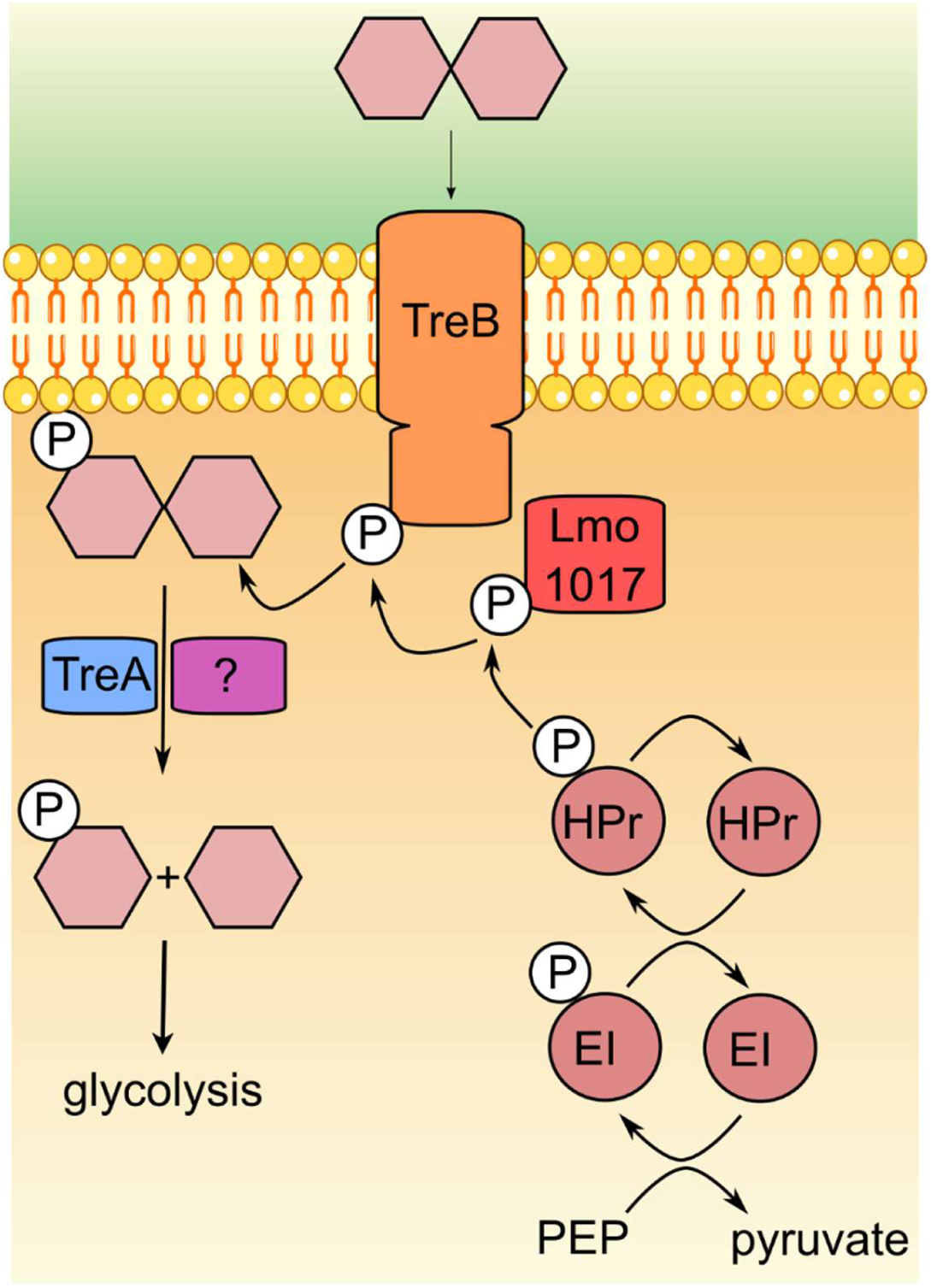
Trehalose import and metabolism. The import and phosphorylation of the disaccharide trehalose is facilitated by the PTS system consisting of the EIIA component Lmo1017 and the EIIBC component TreB. Phosphorylation of Lmo1017 occurs through the canonical PTS phosphorylation cascade, which starts with the phosphorylation of enzyme I (EI) by phosphoenolpyruvate (PEP). The phosphate group is then transferred to the histidine-containing phosphocarrier protein HPr, which subsequently phosphorylates Lmo1017. P∼Lmo1017 transfers the phosphate group to TreB, which in turn phosphorylates trehalose during its import. The resulting trehalose-6-phosphate is hydrolyzed by TreA and an unknown phosphotrehalase and its cleavage products, glucose and glucose-6-phosphate, are then metabolized through glycolysis.

Lack of the phosphotrehalase TreA has been associated with increased tolerance to heat and osmotic stress in *L. monocytogenes* strain 568. This effect is likely due to the accumulation of trehalose, which results from the dephosphorylation of trehalose-6-phosphate (25). Indeed, trehalose can act as a compatible solute in *E. coli*, yeast and other organisms thereby increasing stress hardiness (13, 35). In contrast, accumulation of trehalose-6-phosphate and other phosphorylated sugars are toxic and can reduce the stress tolerance (36–38). While we did not assess stress tolerance of the *treA* mutant, we observed a reduced growth when TreB was overproduced in the *L. monocytogenes* wildtype. This growth defect was not observed when both, TreA and TreB were overexpressed. Overproduction of TreB alone likely leads to enhanced import of trehalose, which cannot be sufficiently hydrolyzed by TreA. The growth defect is thus likely caused by an accumulation of trehalose-6-phosphate.

In *B. subtilis*, the expression of the *treBA* operon is controlled by the repressor TreR (15). We here confirm that TreR, encoded by *lmo1253*, also binds to the *treBA* promoter region and acts as a repressor in *L. monocytogenes*. The *treBA* operon is highly expressed, when *L. monocytogenes* is grown in BHI or LB broth, both of which contain trehalose (25). In contrast, it is barely expressed, when *L. monocytogenes* is grown in LSM glucose. Promoter activity assays further revealed that the expression of the *treBA* operon can still be enhanced in the *treR* deletion strain in response to trehalose, suggesting that the operon might also be controlled by a second regulator. Interestingly, the growth of the *manR* deletion strain was enhanced in LSM trehalose in comparison to the wildtype and similar to the *treR* deletion strain, indicating that ManR could serve as a second regulator of the *treBA* operon.

Even though cellobiose and trehalose only differ in the glycosidic bond that connects two glucose monomers, they exert strong differences in PrfA repression. We thus wondered whether these two sugars are metabolized via different metabolic pathways. Interestingly, ^13^C-labeling experiments revealed that trehalose was not metabolized in the presence of [U-^13^C_6_]glucose, indicating a carbon catabolite repression in *L. monocytogenes*. This regulatory effect has previously been reported for *B. subtilis* and *Streptococcus mutans*, where glucose-1-phosphate inhibited the binding of TreR to the *tre* operator (39). Surprisingly, despite its structural similarity to trehalose, cellobiose does not appear to be subject to catabolite repression in the presence of [U-^13^C_6_]glucose. One possible explanation is that in its saprophytic lifestyle *L. monocytogenes* mainly uses decaying plant material as primary nutrient source (10), and thus also the plant-derived sugar cellobiose. Additionally, the repression of trehalose uptake might also be specifically required to prevent the intracellular accumulation of trehalose-6-phosphate, which has been reported to exert toxic or growth-inhibiting effects in bacteria and fungi (14, 37, 40). No catabolite repression of trehalose metabolism was observed when [U-^13^C_3_]glycerol was supplied as the primary carbon source in LSM. Under these conditions, both trehalose and cellobiose were metabolized, and their carbon was incorporated into identical pathways, including glycolysis, the incomplete TCA cycle, the pentose phosphate pathway, amino acid biosynthesis, and fatty acid biosynthesis. Interestingly, trehalose showed a slower uptake and/or metabolization rate than cellobiose. Further experiments need to be performed to distinguish between these two possibilities.

Due to clear differences in *treBA* expression and the metabolization of trehalose in diverse media, promoter activity assays were performed to assess the repressive effect of trehalose on PrfA. While there was no clear repressive effect when *L. monocytogenes* was grown in BHI amberlite containing trehalose, *P_actA_* activity was reduced in LB amberlite containing trehalose. Since the expression of the *treBA* operon is more then 10-fold higher in LB compared to BHI medium, we speculate that the metabolization rate of trehalose might be enhanced, when *L. monocytogenes* is grown in LB, which ultimately enables trehalose-dependent PrfA repression. The ^13^C-glycerol experiments revealed that trehalose consumption seems to be slower than the consumption of cellobiose, which would explain why the degree of PrfA repression differs between both sugars. Lecithinase activity assays further revealed that the absence of the transcriptional regulators of the glucose- and cellobiose-specific PTS systems ManR and CelR, reduces the repressive effect of glucose and cellobiose, respectively, although no clear growth differences on the respective sugars were observed. It was previously shown that the deletion of *celR* leads to a reduced consumption rate of cellobiose (24). We assume that the glucose consumption rate is also reduced in a *manR* deletion strain and that the decreased transport and/or metabolization of glucose and cellobiose leads to a partial relief of sugar-dependent PrfA repression, which had also been previously observed (23, 24, 31). Therefore, the sugar-dependent repression of PrfA seems to correlate with the transport and/or consumption rate of the corresponding sugar as well as the growth medium used. Altering any of these factors already changes the activity of PrfA and thus the transcription of virulence factors. We therefore propose that sugar-specific transport and/or metabolization rates lead to differences in phosphoenolpyruvate (PEP) availability. Since PEP functions both, as a metabolic intermediate and as a phosphate donor for PTS-dependent transport, variations in its intracellular concentration may act as a regulatory signal that modulates transporter activity as well as virulence through PrfA activity.

## Acknowledgements

We thank Julia Busse for technical assistance and Lea Hahn and Jasmin Petrovac for the help with some experiments. We are grateful to Prof. Jörg Stülke for providing JR and MJ with laboratory space, equipment and consumables and to the Göttingen Center for Molecular Biosciences (GZMB) and the Dorothea Schlözer Programme for financial support. This work was funded by the German research foundation (DFG) grant RI 2920/3-3 to JR and EI 384/16-1 to WE.

