## Supplemental Material for "Trehalose metabolism and its impact on PrfA activity in *Listeria monocytogenes*"

**Supplemental Tables**

**Table S1: Bacterial strains used in this study**

| Unique ID | Strain name and resistance | Source |
|---|---|---|
| ***Escherichia coli* strains** | | |
| ANG124 | DH5α pKSV7; AmpR | (1) |
| ANG4242 | XL1-Blue pIMK2; KanR | (2) |
| ANG5181 | XL1-Blue pPL3e-*lacZ*; CamR | (3) |
| EJR274 | XL1-Blue pPL3e-$P_{plcA}$-$P_{prfA}$-*prfA*; $cat^R$ | This study |
| EJR286 | DH5α pPL3e-$P_{actA}$-*lacZ*; CamR | This study |
| EJR378 | DH5α pKSV7-Δ*celR*; AmpR | This study |
| EJR379 | DH5α pKSV7-Δ*manR*; AmpR | This study |
| EJR382 | DH5α pPL3e-$P_{treBA}$-*lacZ*; CamR | This study |
| EJR383 | DH5α pIMK2-*treBA*; KanR | This study |
| EJR385 | DH5α pIMK2-*treB*; KanR | This study |
| EJR388 | DH5α pKSV7-Δ*treA*; AmpR | This study |
| EJR389 | DH5α pKSV7-Δ*treB*; AmpR | This study |
| EJR390 | DH5α pKSV7-Δ*treR*; AmpR | This study |
| EJR403 | S17-1 pPL3e-$P_{treBA}$-*lacZ*; CamR | This study |
| EJR404 | DH5α pKSV7-Δ*lmo1017*; AmpR | This study |
| EJR409 | DH5α pWH844-*treR*; AmpR | This study |
| EJR415 | DH5α pKSV7-Δ*lmo0862*; AmpR | This study |
| EJR417 | DH5α pIMK2-*lmo0862*; KanR | This study |
| EJR443 | DH5α pKSV7-Δ*lmo0184*; AmpR | This study |
| EJR444 | DH5α pIMK2-*lmo0184*; KanR | This study |
| ***Listeria monocytogenes* strains** | | |
| ANG873 | EGD-e | (4) |

| | | |
|---|---|---|
| BUG2214 | EGD-e Δ*prfA* | (5) |
| LJR334 | EGD-e Δ*prfA* pPL3e-$P_{plcA}$-$P_{prfA}$-*prfA*; ery$^{R}$; short: Δ*prfA* compl. | This study |
| LJR352 | EGD-e pPL3e-$P_{actA}$-*lacZ*; ErmR | This study |
| LJR436 | EGD-e Δ*celR* | This study |
| LJR584 | EGD-e Δ*manR* | This study |
| LJR622 | EGD-e pPL3e-$P_{treBA}$-*lacZ*; ErmR | This study |
| LJR644 | EGD-e pIMK2-*treB*, KanR | This study |
| LJR646 | EGD-e pIMK2-*treBA*, KanR | This study |
| LJR717 | EGD-e Δ*treA* | This study |
| LJR718 | EGD-e Δ*treB* | This study |
| LJR719 | EGD-e Δ*treR* | This study |
| LJR730 | EGD-e Δ*treR* pPL3e-$P_{treBA}$-*lacZ*; ErmR | This study |
| LJR746 | EGD-e Δ*lmo1017* | This study |
| LJR760 | EGD-e pIMK2-*lmo0862*; KanR | This study |
| LJR761 | EGD-e Δ*treA* pIMK2-*lmo0862*; KanR | This study |
| LJR776 | EGD-e pIMK2-*lmo0184*; KanR | This study |
| LJR778 | EGD-e Δ*treA* pIMK2-*lmo0184*; KanR | This study |
| LJR791 | EGD-e Δ*lmo0862* | This study |
| LJR792 | EGD-e Δ*treA* Δ*lmo0862* | This study |
| LJR793 | EGD-e Δ*treA* Δ*lmo0184* | This study |
| LJR794 | EGD-e Δ*lmo0184* | This study |

**Table S2: Primers used in this study**

| Number | Name | Sequence |
|---|---|---|
| JR257 | $P_{plcA}$-$P_{prfA}$-*prfA* rev | ACGCGTCGACCGAATAAAATATAAAACAGTATTCCTC |
| JR312 | $P_{plcA}$-$P_{prfA}$-*prfA* fw | CGGGGTACCGAAATTCGCTTCTAAAGATGAAACG |
| JR434 | *treB* up rev | AGATTCACGATAAGCGCTTGCATCCTTTTTATAGTC |
| JR435 | *treB* down fw | GATGCAAGCGCTTATCGTGAATCTGCAAAAACAACCAAC |
| JR465 | *manR* up fw | AAAGGTACCCCATCCTAATTCATCCCTTTCG |
| JR466 | *manR* up rev | TTGGATTTCATTAAATTCATAAATACGGTCAATACGCTTC |
| JR467 | *manR* down fw | ATTTATGAATTTAATGAAATCCAACCAGAAGTACTTTC |
| JR468 | *manR* down rev | TTTGTCGACCTTCTACGAGGGTTAGTACATC |
| JR471 | *celR* up fw | AAAGGTACCGCGACAAAACTAGAAAATCCGG |
| JR472 | *celR* up rev | ATTGGAGAGCATTTGGAGGACTTCTTCTTTTCTACTA |
| JR473 | *celR* down fw | GAAGTCCTCCAAATGCTCTCCAATCAGGTAGAAG |
| JR474 | *celR* down rev | TTTGTCGACGCTACATGTTGTGGAAATACGG |
| JR489 | $P_{treBA}$ fw | AAAGGATCCCATTTTCAACAGCGCCACCC |
| JR490 | $P_{treBA}$ rev | TTTGTCGACATAGTCAACCATAAAATTTCCTCCTTTTTAAC |
| JR491 | pIMK2-*treBA* fw | AAACCATGGTTGACTATAAAAAGGATGCAAGCG |
| JR492 | pIMK2-*treBA* rev | TTTGTCGACTTATTGTATTAATGTTAATGTTTCATATGGCG |
| JR493 | pIMK2-*treB* rev | TTTGTCGACTTAGTTGGTTGTTTTTGCAGATTCAC |
| JR494 | *treA* up fw | AAAGGTACCGGTGTTTATAGTTTCCTTTGGCTC |
| JR495 | *treA* up rev | TGTTTCATATGGATAAATTGTTTTTTGAGCAAACTTTGTCAT |
| JR496 | *treA* down fw | AAAACAATTTATCCATATGAAACATTAACATTAATACAATAA |
| JR497 | *treA* down rev | TTTGTCGACCTAACAAATTTCGGTTACAATTAGGG |
| JR501 | *treR* up rev | ATCAATAAAACGAATGTCAAAAAACTTATTTTTCTTATTCAAATTT |
| JR502 | *treR* down fw | TTTTTTGACATTCGTTTTATTGATTTTGCTAGAAGACG |
| JR507 | *treB* up fw | AAAGGTACCCCGTTCTGCCCAAACTGTTC |

| | | |
|---|---|---|
| JR508 | *treB* down rev | TTTGTCGACGTACATTCGGATTGGCCCAG |
| JR511 | *treR* up fw | AAAGGTACCCCTCTAAATAGGCACTCTGGAG |
| JR512 | *treR* down rev | TTTGTCGACCGGACTTAGTGAGCAGGAAG |
| JR551 | pIMK2-*lmo0862* fw | AAAGGATCCCGAGTTTTGGCGTCGTAGTGTG |
| JR552 | pIMK2-*lmo0862* rev | TTTGTCGACTTATTCCTCCACTCTAAGCGC |
| JR561 | pIMK2-*lmo0184* fw | AAAGGATCCCAAAGAAAAAGATTGGTGGAAAAAAAGC |
| JR562 | pIMK2-*lmo0184* rev | TTTGTCGACTTACTTGATTTGATAAACGATTGCTTC |
| JR563 | *lmo0184* up fw | AAAGGTACCGGGCTTTTGGTGCGGACG |
| JR564 | *lmo0184* up rev | GATTGCTTCGTAGCTTTTTTTCCACCAATCTTTTTCTTTC |
| JR565 | *lmo0184* down fw | TGGAAAAAAAGCTACGAAGCAATCGTTTATCAAATCAAG |
| JR566 | *lmo0184* down rev | TTTGTCGACGCATAATTCCGCCTACTTCTTC |
| JS21 | *lmo0862* up fw | AAAGGATCCCGCTATATACTGACAACATTACTC |
| JS22 | *lmo0862*up rev | GTTCCCCTGAAGAAACACACTACGACGCCAAAAC |
| JS23 | *lmo0862*down fw | CGTAGTGTGTTTCTTCAGGGGAACTATCAGCTT |
| JS24 | *lmo0862*down rev | TTTGTCGACCGATGTGACCAAGCACGC |
| NW11 | *lmo1017* up fw | AAAGGTACCCAATCCAGATAAAGTAGCAAAATGGA |
| NW12 | *lmo1017* up rev | GTTTTCCCTGCTTCCGCCATGCATCTTCCTCCTAGCAAT |
| NW13 | *lmo1017* down fw | CTAGGAGGAAGATGCATGGCGGAAGCAGGGAAAACC |
| NW14 | *lmo1017* down rev | AAAGTCGACGGTAACTCGCTTCGATATCTTTG |
| NW21 | pWH844-*treR* fw | AAAGGATCCTTGAATAAGAAAAATAAGTTTTTTGACATTTATTTAG |
| NW22 | pWH844-*treR* rev | AAAGTCGACTTAGCGTCTTCTAGCAAAATCAATAAAAC |
| NW23 | EMSA $P_{treBA}$ fw | ATAGTCAACCATAAAATTTCCTCCTTTTTAAC |
| NW24 | EMSA $P_{treBA}$ rev | CATTTTCAACAGCGCCACCC |
| NW25 | EMSA $P_{cadA}$ fw | CGTACAAGACATACCGTCTAC |
| NW26 | EMSA $P_{cadA}$ rev | CATTTACTTACCTTAAGACAAG |

**Supplemental Figures**

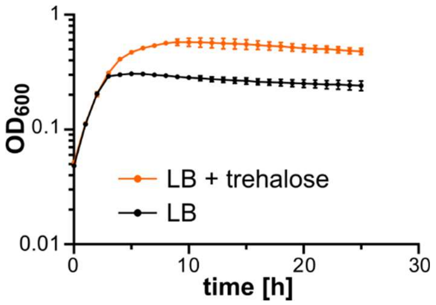


**Fig. S1: Enhanced growth of *L. monocytogenes* in the presence of trehalose.** Overnight cultures of the *L. monocytogenes* wildtype strain EGD-e were diluted to an $OD_{600}$ of 0.1 in fresh LB medium. When the cultures reached an $OD_{600}$, cells were collected, washed twice in LB medium or LB medium containing 25 mM trehalose and $OD_{600}$ adjusted as described in the methods section. The growth was monitored for 25 hours using a plate reader. The average values and standard deviations of the $OD_{600}$ readings of three independent experiments were plotted.

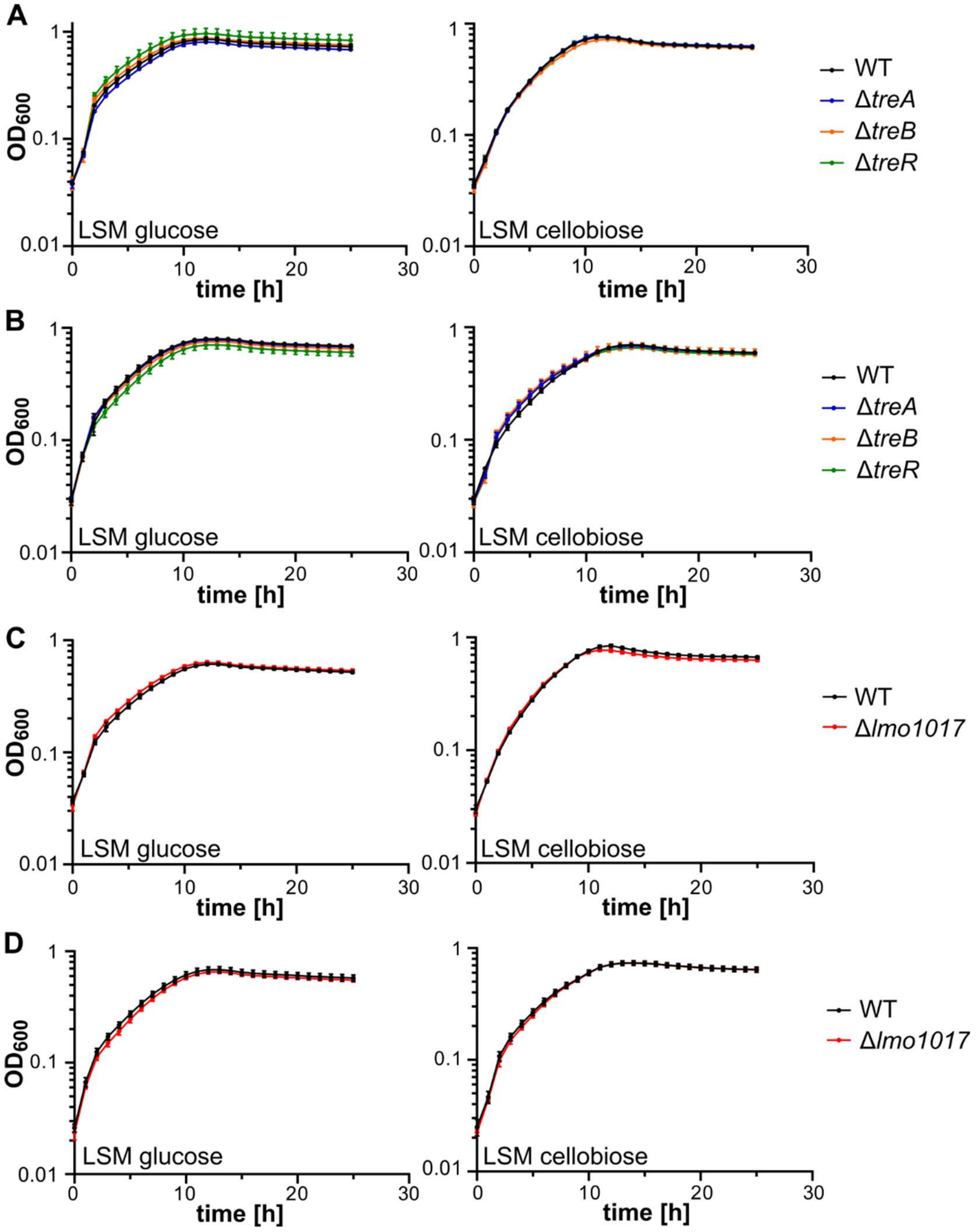


**Fig. S2: Growth of *treA*, *treB*, *treR* and *lmo1017* deletion strains on LSM glucose and cellobiose.** (A, C) The indicated *L. monocytogenes* strains were grown overnight in BHI broth, diluted to an $OD_{600}$ of 0.1 in fresh BHI broth and grown until an $OD_{600}$ of 0.3 was reached. (B, D) *L. monocytogenes* strains were grown overnight in LSM glucose, diluted to an $OD_{600}$ of 0.1 in fresh LSM glucose and grown until an $OD_{600}$ of 0.3 was reached. (A-D) Cells were collected, washed and $OD_{600}$ adjusted as described in the methods section. The growth was monitored for 25 hours using a plate reader. The average values and standard deviations of the $OD_{600}$ readings of three independent experiments were plotted.

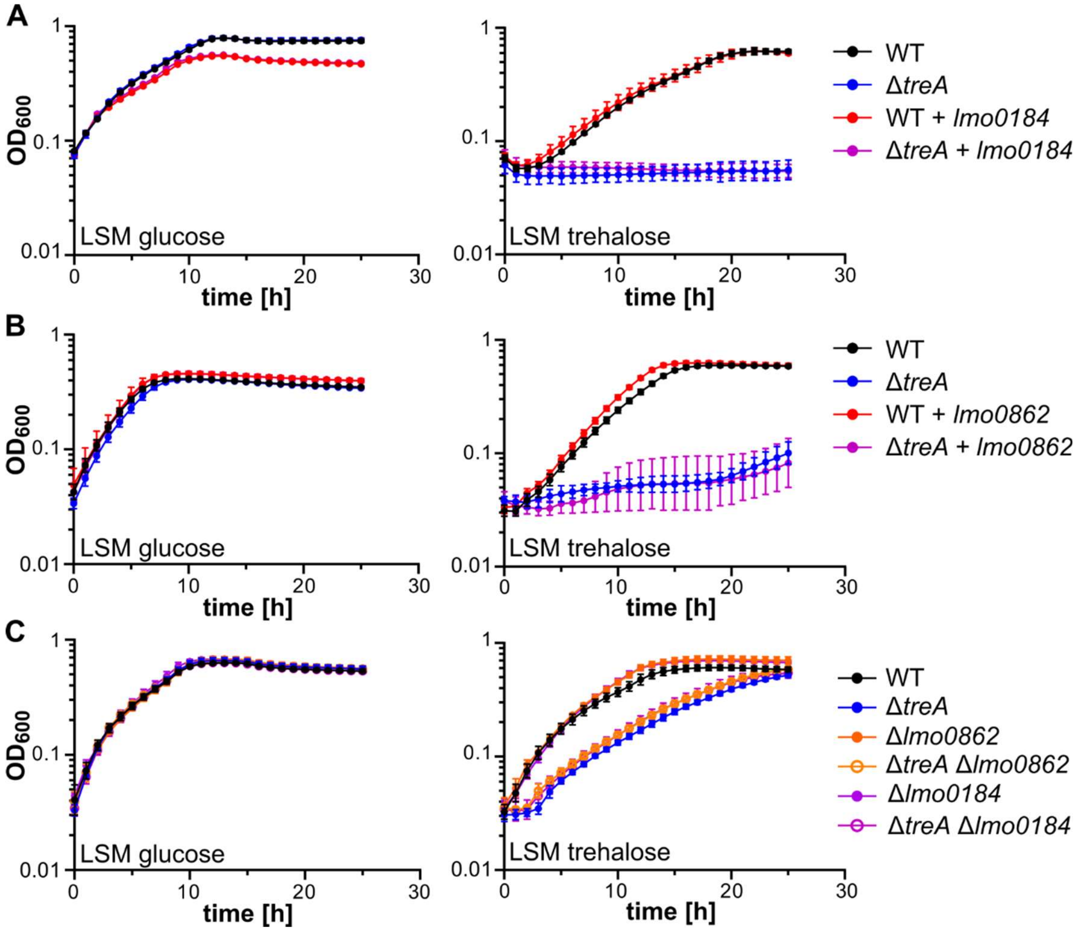


**Fig. S3: Lmo0184 und Lmo0862 are not required for trehalose metabolism.** (A-B) *L. monocytogenes* strains were grown overnight in LSM glucose, diluted to an $OD_{600}$ of 0.1 in fresh LSM glucose and grown until an $OD_{600}$ of 0.3 was reached. (C) The indicated *L. monocytogenes* strains were grown overnight in BHI broth, diluted to an $OD_{600}$ of 0.1 in fresh BHI broth and grown until an $OD_{600}$ of 0.3 was reached. (A-C) Cells were collected, washed and $OD_{600}$ adjusted as described in the methods section. The growth was monitored for 25 hours using a plate reader. The average values and standard deviations of the $OD_{600}$ readings of three independent experiments were plotted.

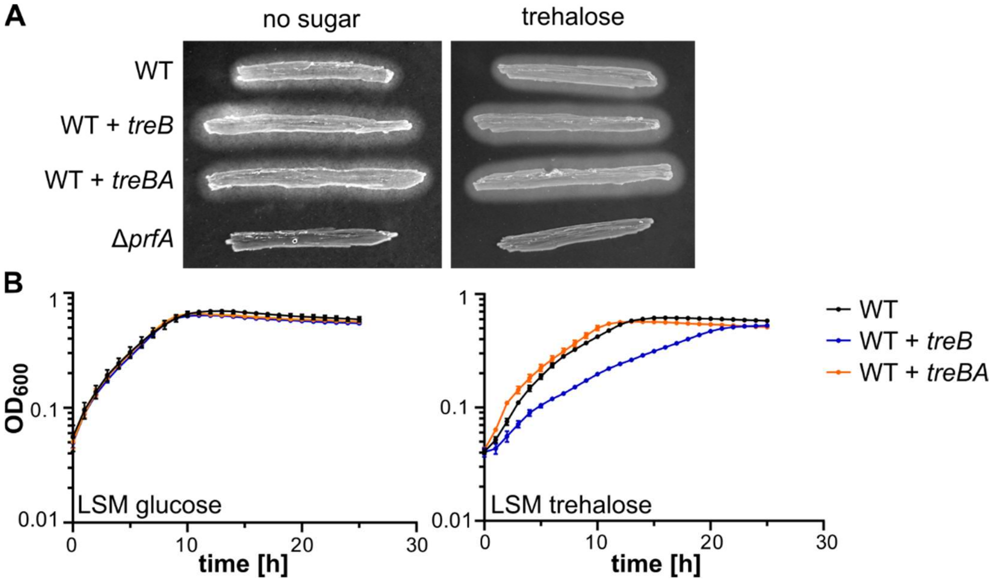


**Fig. S4: Overexpression of *treB* leads to growth defect in LSM trehalose.** (A) PlcB activity of the *L. monocytogenes* wildtype strain EGD-e (WT), the *treB* overexpression strain (LJR644; WT + *treB*), the *treBA* overexpression strain (LJR646; WT + *treBA*) and the *prfA* mutant was tested on LB agar plates containing activated charcoal and egg yolk or on plates that additionally contained 25 mM trehalose. (B) Overnight cultures of the *L. monocytogenes* wildtype strain EGD-e, the *treB* overexpression strain and the *treBA* overexpression strain were diluted to an $OD_{600}$ of 0.1 in fresh BHI medium. When the cultures reached an $OD_{600}$, cells were collected, washed and $OD_{600}$ adjusted as described in the methods section. The growth in LSM glucose and LSM trehalose was monitored for 25 hours using a plate reader. The average values and standard deviations of the $OD_{600}$ readings of three independent experiments were plotted.

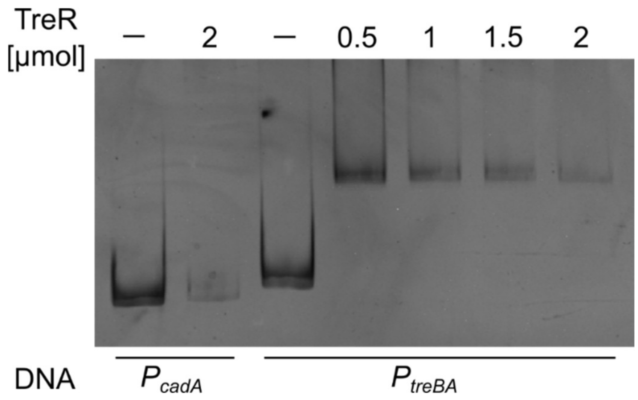


**Fig. S5: TreR binds to the $P_{treBA}$ promoter region.** Increasing concentrations of recombinant His-TreR were incubated with a 236-bp fragment containing the *treBA* promoter (lanes 4-7). A reaction of 2 µmol TreR with a 200-bp fragment containing the *cadA* promoter was used as negative control (lane 2). Reactions without protein were used as additional control (-).

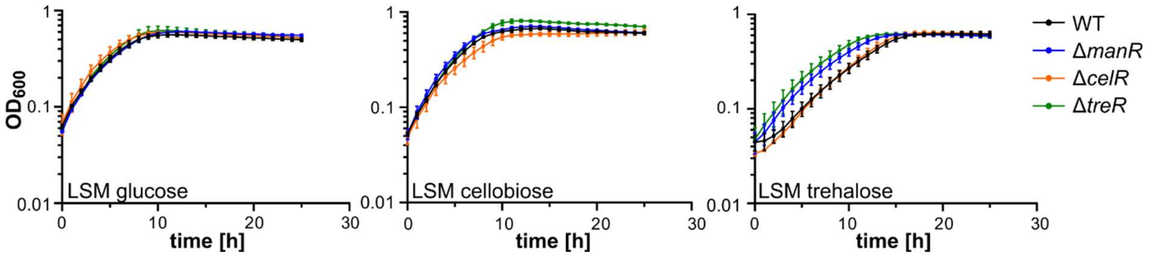


**Fig. S6: Growth of *manR*, *celR* and *treR* mutants.** *L. monocytogenes* strains were grown overnight in LSM glucose, diluted to an $OD_{600}$ of 0.1 in fresh LSM glucose and grown until an $OD_{600}$ of 0.3 was reached. Cells were collected, washed and $OD_{600}$ adjusted as described in the methods section. The growth was monitored for 25 hours using a plate reader. The average values and standard deviations of the $OD_{600}$ readings of three independent experiments were plotted.
